# The genetic architecture of multiple mutualisms and mating system in *Turnera ulmifolia*

**DOI:** 10.1101/2022.03.31.484762

**Authors:** Jason R. Laurich, Christopher G. Reid, Caroline Biel, Tianbu Wu, Christopher Knox, Megan E. Frederickson

**Affiliations:** Department of Ecology and Evolutionary Biology, University of Toronto, M5S 3B2, Toronto, Canada; Faculty of the Environment, Simon Fraser University, V5A 1S6 Burnaby, Canada

**Author notes:** Corresponding Author: Jason R. Laurich, 1(.

**Keywords:** multiple mutualisms, pollination, biotic defence, seed dispersal, G matrix

## Abstract

Plants often associate with multiple arthropod mutualists. These partners provide important services to their hosts, but multiple interactions can constrain a plant’s ability to respond to complex, multivariate selection. Here, we quantified patterns of genetic variance and covariance among rewards for pollination, biotic defence, and seed dispersal mutualisms in multiple populations of *Turnera ulmifolia* to better understand how the genetic architecture of multiple mutualisms might influence their evolution. We phenotyped plants cultivated from 17 Jamaican populations for several mutualism and mating system-related traits. We then fit genetic variance-covariance (G) matrices for the island metapopulation and the 5 largest individual populations. At the metapopulation level, we observed significant positive genetic correlations among stigma-anther separation, floral nectar production, and extrafloral nectar production. These correlations have the potential to significantly constrain or facilitate the evolution of multiple mutualisms in *T. ulmifolia* and suggest that pollination, seed dispersal, and dispersal mutualisms do not evolve independently. In particular, we found that positive genetic correlations between floral and extrafloral nectar production may help explain their stable co-existence in the face of physiological trade-offs and negative interactions between pollinators and ant bodyguards. Locally, we found only small differences in G among our *T. ulmifolia* populations, suggesting that geographic variation in G may not shape the evolution of multiple mutualisms.

## Introduction

Organisms typically interact with multiple mutualists and antagonists, either simultaneously or sequentially (Strauss 2014; TerHorst et al. 2018). The outcomes of these multiple interactions are often difficult to predict (Strauss 2014), given the potential for different community members to exert opposing fitness effects (e.g., Sletvold et al. 2015) or to interact non-additively (e.g., Afkhami et al. 2014; Afkhami and Stinchcombe 2016). Incorporating the multiple interactions that capture the complexity of real communities has improved our understanding of the ecology of species interactions (Strauss 2014; TerHorst et al. 2018), but relatively few studies have explored how populations adapt to the multiple species with which they interact.

The translation of ecological complexity into the evolution of species interactions is complicated by the fact that organisms do not respond to selection imposed by individual partners in isolation. Rather, selection is a multivariate process acting on many traits, and its effects depend on genetic features such as pleiotropy and linkage disequilibrium among loci targeted by selection (Lande 1979; Kopp and Matuszewski 2014; TerHorst et al. 2018). These genetic features may result in genetic covariances among traits that affect interactions with different community members, and can either facilitate or constrain phenotypic evolution, depending on whether the principal axes of co-variation are aligned with those of selection (Arnold 1992; Agrawal and Stinchcombe 2009; Wise and Rausher 2013; Teplitsky et al. 2014; TerHorst et al. 2018; Assis et al. 2020). More formally, Lande’s equation, *R* = *Gβ*, predicts that the vector of responses to selection on multiple traits, *R*, equals the product of the genetic variance-covariance matrix, *G*, and the vector of selection gradients acting on those traits, *β* (Lande and Arnold, 1983; Hansen and Houle, 2008). Traits that mediate species interactions will therefore have a correlated response to selection whenever they are genetically correlated, such that multiple species interactions may evolve in tandem. Here, we estimated genetic covariances among multiple traits that mediate plant mutualisms with ants and pollinators, and simulated selection (i.e, *β*) to understand how the genetic architecture (i.e., G) of these traits may affect their evolution.

While genetic covariances may in general have only limited effects on rates of adaptation (Agrawal and Stinchcombe 2009), the evolution of traits involved in multiple species interactions may be especially susceptible to this form of constraint or facilitation. Multiple species interactions are often mediated by several traits in a focal species in ways that can lead to correlational selection and the emergence of genetic correlations among them (e.g., Jones et al. 2004, 2012; Penna et al. 2017; Svensson et al. 2021). Correlational selection occurs when fitness depends on interactive effects of two or more traits and is known to influence G-matrix evolution (Svensson et al., 2021). Correlational selection is likely a common outcome of multiple species interactions (e.g., Wise and Rausher, 2013), because of widespread context-dependency in ecological interactions (Chamberlain et al., 2014). Thus, ecological interactions among multiple mutualists could lead to strong genetic correlations among the multiple traits mediating these interactions (Jones et al. 2012; Guimarães *et al*. 2017; Penna et al. 2017). Predicting patterns of correlational selection in multiple species interactions is, however, complicated by the ecological complexity of real communities. When multiple partner species interact and affect the fitness landscape of multiple interactions (Afkhami et al. 2014), they fundamentally change the nature of multivariate selection, creating new patterns of correlational selection dependent on ecological context (e.g., Wise 2010; Wise and Rausher 2013; Friman and Buckling 2013; Wise and Rausher 2016; Guimarães *et al*. 2017; Keith and Mitchell-Olds 2019; Afkhami et al. 2021). Moreover, phenotypes that affect more than one ecological interaction (either directly or through diffuse interactions among partner species) also create the possibility of antagonistic or synergistic effects of multiple species interactions on the responses of even single traits to selection (Cuautle et al. 2005; Sletvold et al. 2015; Wise and Rausher 2016; Villamil et al. 2019; Afkhami et al. 2021).

A growing body of research has begun to shed light on how organisms evolve in response to multiple antagonisms, but very few studies have investigated trait evolution in the context of multiple mutualisms (Guimarães *et al*., 2017). Multiple mutualisms are ubiquitous; many organisms interact with several partners, often from different guilds (Stanton, 2003; Dáttilo et al., 2016; Guimarães *et al*., 2017). Plants associate with many beneficial organisms, typically including some combination of animal pollinators (Ollerton et al., 2011), seed dispersers, and biotic defenders such as ant bodyguards (Marazzi et al., 2013; Weber and Keeler, 2013) in addition to mycorrhizal fungi and diverse microbiomes (moreover, microbiomes themselves are often a type of multiple mutualism). Here, we focus on the aboveground mutualistic interactions of *Turnera ulmifolia* (Passifloraceae), a weedy neotropical plant that associates with a diversity of arthropod mutualists, including pollinators and both defensive and seed-dispersing ants (Barrett and Shore, 1987; Dutton et al., 2016a,b). In *T. ulmifolia,* these interactions are mediated by multiple plant traits, principally floral nectar that is consumed by insect pollinators, and elaiosomes (fleshy, lipid-rich appendages attached to seeds) and extrafloral nectar that are consumed by ants that disperse seeds and defend the plant against herbivores (Barrett and Shore, 1987; Cuautle et al., 2005; Dutton et al., 2016a,b; Villamil et al., 2019).

Conflict between ants and pollinators, which consume similar nectar rewards (extrafloral and floral nectar) that are often in close proximity on plants (Martínez-Bauer *et al*., 2015; Dutton et al., 2016a; Keith and Mitchell-Olds, 2019; Villamil et al., 2019), can give rise to non-additive effects of pollinators and ants on plants (i.e., multiple mutualist effects, Afkhami et al., 2014). Direct interactions between ants and pollinators are common, as aggressive ant mutualists can deter pollinators from visiting in addition to reducing herbivory (Ness, 2006; Cembrowski et al., 2014; Villamil et al., 2018). Biotic defense and pollination are further linked through herbivores, which can be attracted to showy floral displays. This can create an ecological trade-off between attracting mutualists and repelling antagonists (Knauer and Schiestl, 2017), such that the evolution of floral traits affecting pollination may be driven not only by pollinators, but also by herbivores and the effectiveness of plant defenses against them (Strauss et al., 1996; Ågren et al., 2013; Kessler et al., 2015; Sletvold et al., 2015; Soper Gorden and Adler, 2016; Thompson and Johnson, 2016; Gervasi and Schiestl, 2017; Ramos and Schiestl, 2019; Cabin et al., 2022). We might therefore expect to see correlational selection favoring some trait combinations (e.g., greater floral rewards together with high investment in anti-herbivore defense), potentially leading to genetic covariances among plant traits. Furthermore, the existence of genetic covariances, such as that suggested by the genetic and physiological links between the production of biochemically similar rewards like extrafloral and floral nectar (Heil 2011; Dutton et al. 2016a), may feed back to affect the evolution of mutualistic plant traits.

Multiple mutualisms may also contribute to the coordinated evolution of multivariate phenotypes characteristic of evolutionary syndromes. For example, the evolution of selfing in plants is often described as a ‘syndrome’ that involves a set of linked phenotypes, including reduced floral size, nectar volume, and stigma-anther separation (e.g., Belaoussoff and Shore 1995; Sicard and Lenhard 2011). We expect selfing to also co-vary with seed dispersal and anti-herbivore defense traits in plants, which can both be mediated by mutualisms. Selfing is often a mechanism of reproductive assurance in good colonizers (Baker, 1955; Pannell and Barrett, 1998) and if natural selection favours selfing in highly dispersive genotypes, these traits may co-vary genetically. Similarly, in the small, isolated populations established by good dispersers at range edges or in ephemeral habitats, plants have the potential to escape their predators and parasites and may experience reduced selection to maintain specialized, constitutive defenses, sometimes in favour of phenotypically plastic, inducible defenses (Chevin and Lande 2011; Campbell and Kessler 2013; Brudvig et al. 2015). Over time, this may lead genetic covariances between dispersal and defense traits to build up. In our study species, we have further reason to think that dispersal and anti-herbivore defense traits may be linked; *T. ulmifolia*’s seeds are dispersed by ants that are attracted not only to elaiosomes, but to extrafloral nectar, which also has a defensive function (Cuautle et al. 2005; Dutton et al. 2016b).

Ecological and evolutionary links among reproduction, defense, and dispersal in plants raise the possibility that traits governing the multiple mutualisms mediating these processes do not evolve independently. Indeed, similarity in the composition of plant rewards like floral and extrafloral nectar suggests a shared genetic architecture (Afkhami and Stinchcombe, 2016; Dutton et al., 2016a), and previous studies of the evolution of biotic defense and pollination at broad phylogenetic scales have hinted at correlated trait evolution (Chamberlain and Rudgers, 2012). Whether plant traits associated with multiple mutualisms are genetically correlated to one another is however not adequately understood, and traits that intuitively seem linked may not always be (Ossler and Heath, 2018). Furthermore, while phenotypic covariances between ecologically important traits are common, these can be environmentally induced, rather than genetic, and parsing genetic from environmental covariances requires an appropriate study design (Stinchcombe et al., 2002). More generally, evolutionary ecology requires a better understanding of how the genetic architecture of these traits interact with their complex ecology to determine the outcome and evolutionary trajectories of multiple mutualisms (Lau and Terhorst, 2015; Guimarães *et al*., 2017; TerHorst et al., 2018; Assis et al., 2020).

Here, we estimate genetic variance-covariance (G) matrices for phenotypic traits associated with multiple mutualisms in *T. ulmifolia*. We grew plants in a common greenhouse environment and measured plant investment in rewards to pollinators, bodyguards, and seed dispersers, as well as variation in mating system, and fit G-matrices for the 5 largest populations we sampled as well as across the island of Jamaica. We fit both local and island-wide G-matrices to assess trait covariances at different spatial scales; *T. ulmifolia* is a good colonizer that readily spreads to new regions, suggesting few barriers to gene flow at small spatial scales. Because we grew plants from seeds of known parentage in a common environment, we could estimate the total genetic variance for each trait as well as genetic covariances between each pair of traits without the confounding effects of environmental variation. Specifically, we addressed the following questions: (1) Does the production of floral nectar (FN), extrafloral nectar (EFN), and elaiosomes covary genetically in *T. ulmifolia*, and do these mutualistic traits covary with stigma-anther separation? (2) How variable are the G matrices of different *T. ulmifolia* populations across a small geographic range? and (3) How might G facilitate or constrain the evolution of multiple mutualisms under simulated, but realistic selection scenarios?

## Materials and Methods

### Study system

*Turnera ulmifolia* is a continuously flowering, ruderal, herbaceous perennial that readily colonizes roadsides and disturbed habitats in its native range of the Caribbean (Barrett, 1978; Barrett and Shore, 1987). Formerly identified as *T. ulmifolia* var. *angustifolia* within the *Turnera ulmifolia* L. complex characterized by extensive chromosomal variation (Urban, 1883; Shore and Barrett, 1985), all known populations of *T. ulmifolia* are hexaploid (López *et al*., 2013). *Turnera ulmifolia* is self-compatible and its populations are generally small, ephemeral, and moderately inbred (Barrett and Shore, 1987; Belaoussoff and Shore, 1995; Dutton et al., 2016a). Though self-compatible, its flowers are conspicuous (figure 1) and are visited by pollinating insects and hummingbirds that increase seed set (Barrett, 1978; Barrett and Shore, 1987; Cuautle and Rico-Gray, 2003). These flowers are ephemeral, opening in the morning around 7 am and then closing and wilting in the early afternoon (Barrett, 1978). *Turnera ulmifolia*’s out-crossing rate is correlated with stigma-anther separation, a variable trait in Jamaica (our study location) that suggests directional selection for increased out-crossing in some populations (Barrett and Shore, 1987; Shore and Barrett, 1990; Belaoussoff and Shore, 1995).

**Figure 1:**
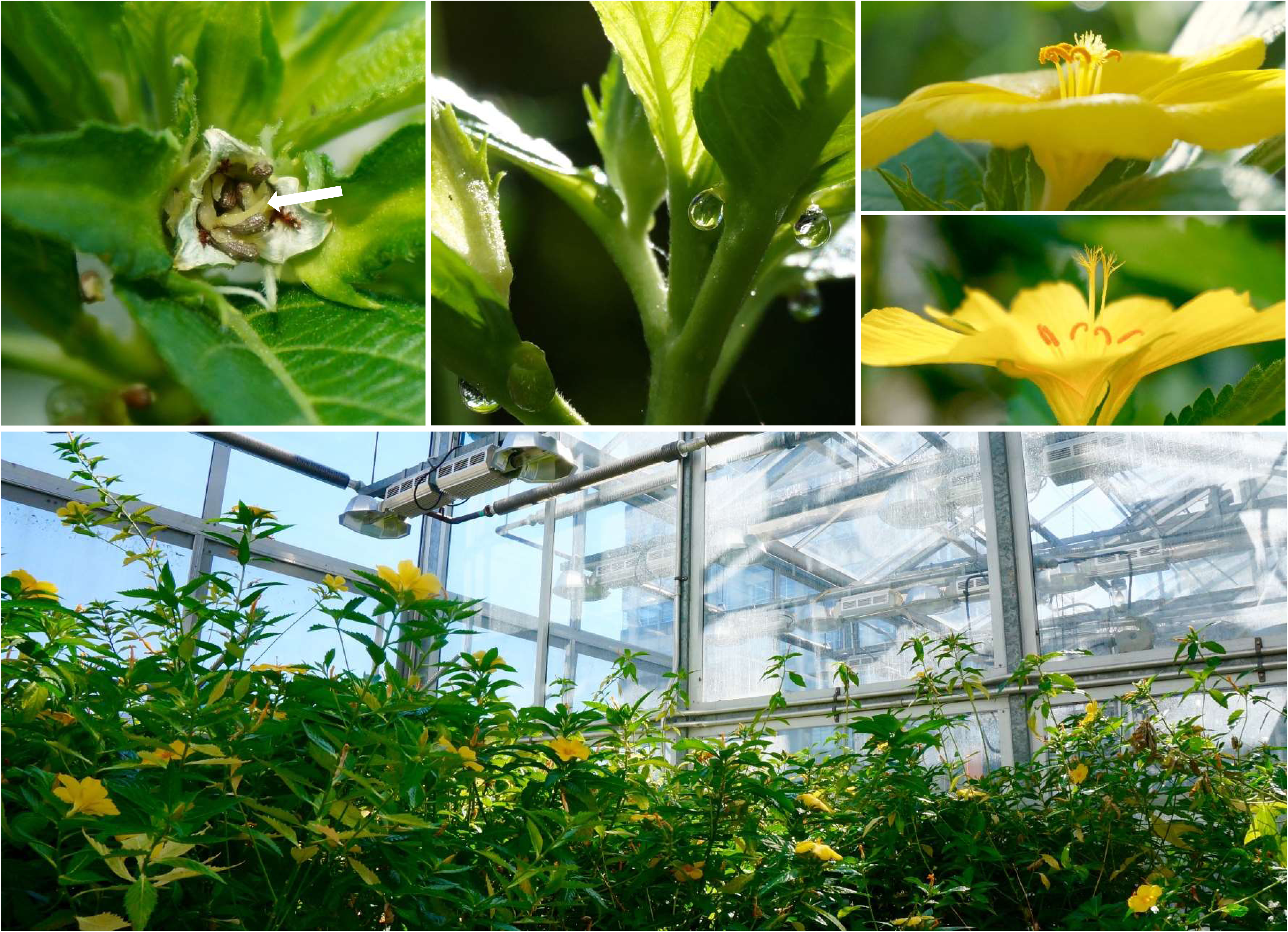
Photographs of *Turnera ulmifolia*. Top row, left: elaiosomes (indicated by white arrow) are visible as fleshy, beige structures affixed to brown seeds within a dehisced capsule. Top row, centre: extrafloral nectar (EFN) accumulating at nectaries on the petioles of leaves. Top row, right: flowers, showing variation in stigma-anther separation. Bottom: plants growing in the greenhouse. Photographs by C. Reid.

In addition to producing large, nectar-rich flowers, *T. ulmifolia* produces extrafloral nectar (EFN) from glands located on the petioles of leaves (Dutton et al., 2016a, figure 1). This EFN attracts ant and wasp bodyguards that defend plants against herbivores and increase reproductive success (Cuautle and Rico-Gray, 2003; Dutton et al., 2016a). *Turnera ulmifolia*’s seeds are dispersed by mutualistic ants which collect seeds from dehisced capsules and feed the attached lipid-rich appendages (elaiosomes, figure 1) to their larvae. The recruitment of seed-dispersing ants to *T. ulmifolia*’s seed capsules is also affected by the production of EFN, which they collect and consume (Salazar-Rojas et al., 2012; Dutton et al., 2016b). Prior to this study, little was known about variation in mutualistic traits other than stigma-anther separation in *T. ulmifolia*, and very little is currently known about the extent of gene flow among Jamaican *T. ulmifolia* populations or the relevant scale at which selection acts on this species.

### Plant collection and cultivation

In 2016, we collected seeds from 218 wild plants distributed across 17 populations of *T. ulmifolia* in Jamaica (figure 2). At each site, we collected seed from up to 20 maternal plants, though only six of the populations (Brown’s Town, Cave, Haining, Mosquito Cove, Murdock, and Saint Ann’s Bay) comprised 20 or more mature plants (table A1). We germinated seeds in a 1:1:1:1 mix of topsoil, peat loam, manure, and concrete sand and transplanted seedlings into 3.5” pots in the greenhouse at the University of Toronto (15L:9D cycle). We watered plants daily and fertilized them bi-weekly. The greenhouse temperatures were set to 20-24 °C and 18-19 °C during the day and night, respectively, although the temperatures inside varied somewhat with season.

**Figure 2:**
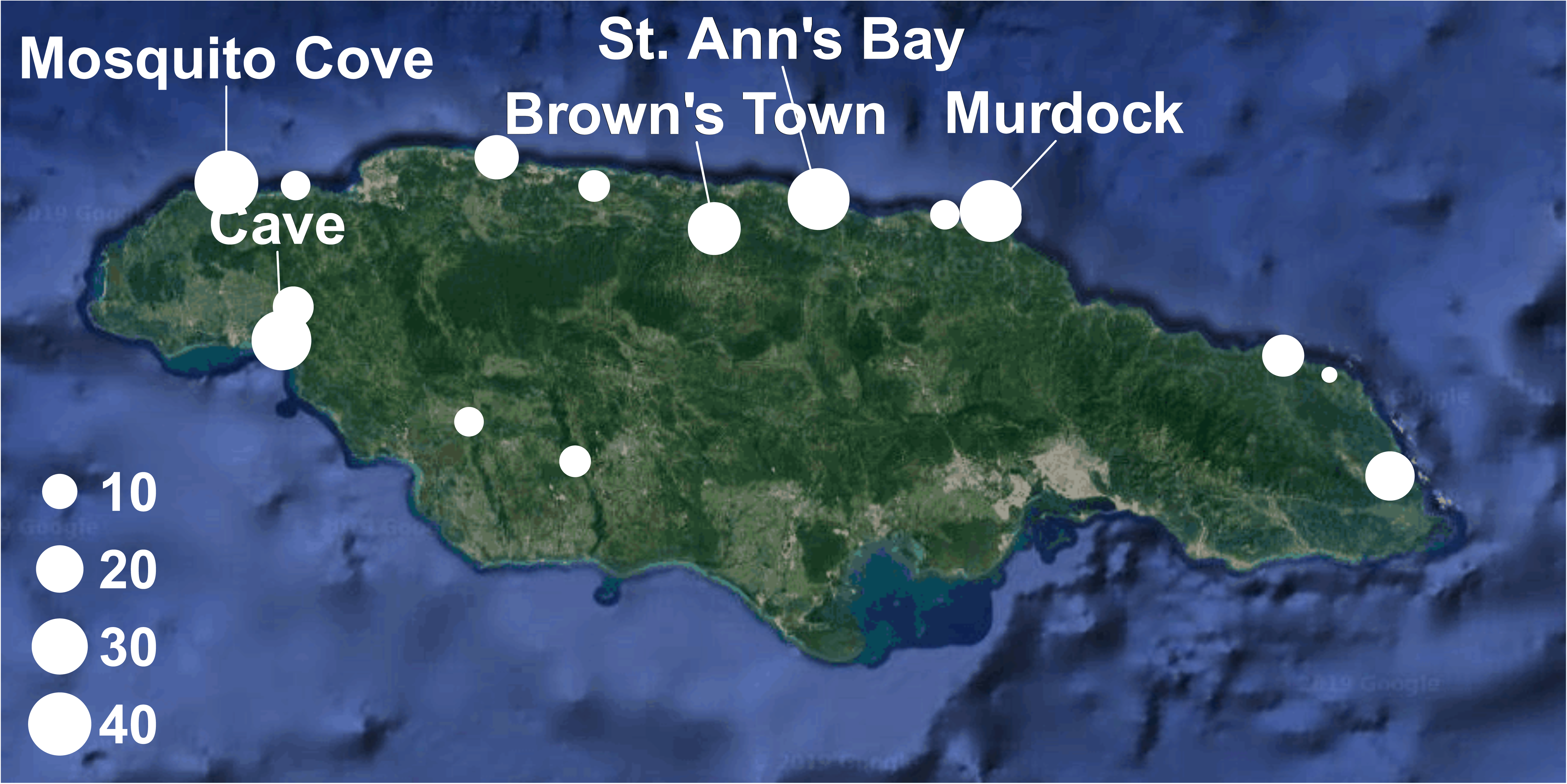
Map of *Turnera ulmifolia* collection sites in Jamaica. Population size is represented by circle diameter and the five well-sampled populations where we collected seeds from more than 20 maternal plants are labelled.

The majority of plants reached reproductive maturity after approximately 8 months. To reduce maternal environmental effects, we generated selfed seed by hand-pollinating flowers from each plant with pollen from the same flower and collected mature seeds after approximately 20-25 days (Dutton et al., 2016a). In March 2017, we germinated seed from 197 maternal families (at most one per plant from which we collected seeds in the field) and planted 10 seedlings per maternal line, for a total of 1970 plants. Each seedling was planted in a 5” pot. We randomly assigned plants to two greenhouses and continued to care for them as described above.

### Trait measurements

Once plants began flowering, we measured stigma-anther separation using calipers. To quantify plant investment in pollination, defense, and seed dispersal mutualisms, we measured floral nectar (FN) production, EFN production, and elaiosome mass, respectively (see figure 1). As the ephemeral flowers of *T. ulmifolia* are a highly dynamic and variable resource for pollinators, we quantified floral resources at the pollinator-relevant level of individual flowers. We used microcapillaries to collect FN from a single open flower per plant. The number of extrafloral nectaries are substantially more stable over time, with new nectaries being produced only with the growth of new leaves. Furthermore, given that ants forage and collect EFN from the entirety of individual plants in the field, we measured the abundance of nectar rewards available for biotic defenders at the plant level. Twenty-four hours prior to collecting EFN, we thoroughly washed plants with water to dislodge any nectar that had previously accumulated or crystallized on extrafloral nectaries. The next day, we collected all the EFN that had accumulated on the plants’ active nectaries in 24 hours using microcapillaries; we later divided by the number of nectaries to get a measure of EFN production per nectary. To determine the total sugar content of FN and EFN in each sample, we diluted samples with known volumes of distilled water and measured total volume with calipers. We measured the sugar concentration of FN and EFN in degrees Brix using a hand-held refractometer. We then calculated the total sugar mass of the original samples (see Appendix B for full details). Finally, we collected autogamous seed from mature seed capsules and dried them for several months at room temperature. We separated intact elaiosomes from three seeds per plant and weighed them on a microbalance. Some plants died or failed to mature; in total we were able to collect a full dataset of all mutualistic phenotypes from 1410 plants across 193 maternal families.

### Data analysis

We performed all statistical analyses in R version 3.5.1 (R Core Team, 2018). Briefly, we fit a common genetic variance-covariance matrix (G_W_, Chenoweth et al., 2010) for the Jamaican metapopulation of *T. ulmifolia* using Bayesian methods. Because the extent of gene flow among populations and the scale at which natural selection acts on *T. ulmifolia* is unknown, we also estimated G for our five largest individual populations, and evaluated the extent to which they differed from one another using three different approaches. We also fit a D matrix summarizing multivariate divergence in trait means among our multiple populations, and assessed whether it is aligned with our island-wide estimate of G_W_ to determine whether G has constrained the direction of phenotypic divergence in *T. ulmifolia*. Finally, we used our local estimates of G to assess evolutionary constraint and facilitation in the multiple mutualisms of *T. ulmifolia* and calculated heritabilities for elaiosome mass, EFN production, FN production, and stigma-anther separation.

#### Estimation and significance testing of G matrices

We estimated the common genetic variance-covariance (G_W_) matrix for the island metapopulation using all plants for which we had all trait data (1410 plants from 193 maternal inbred families). G matrices for this metapopulation, and those later fit for individual populations, were estimated through the implementation of multivariate “animal models” that included elaiosome mass, EFN production, FN production, and stigma-anther separation as response variables. In these models, a vector of individual phenotypes (*Y*) is fitted as a function of the mean phenotypes of a population (*µ*, fixed effects), the genetic breeding values of individuals (*g*, random effects) and residual error (*e*). A fuller description of the multivariate animal model we implemented is:

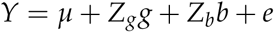

where *Y* is the vector of standardized phenotypes for all plants and *µ* is the vector of mean phenotypes. The genetic effects are represented by the vector *g* and the associated design matrix *Z_g_*, while the random effects of block are represented by the vector of block effects *b* and its corresponding design matrix, *Z_b_* (Kruuk, 2004; Teplitsky et al., 2014). Variance and covariances among the estimated breeding values of individuals within a population (*g*) then yield the genetic variance-covariance matrix.

We fit our models with the ‘MCMCglmm’ package in R (Hadfield, 2010; R Core Team, 2018). We log-transformed total floral nectar sugar mass (FN), and standardized all phenotypic data to a mean of 0 and a variance of 1 against the grand mean and variance of the Jamaican metapopulation. We standardized trait data to the island-wide mean and variance, not within each population separately, in order to estimate G_W_ as if the whole island of Jamaica were a single *T. ulmifolia* population not subject to limited dispersal (which it may be); for the same reason, we did not include population as a random effect when estimating G_W_. For comparison, we also standardized the phenotypic data to a mean of 0 but not to unit variance and re-fit the island-wide G-matrix, G_W_ (but results were similar, see table A11). We applied a set of weakly informative inverse Wishart priors that differed in their specification of predicted genetic variance (*V* = 0.25, 0.333, and 0.5) and degree of belief (*nu* = *n* - 1, *n*, and *n* + 1, where *n* is the number of traits). All priors produced similar G matrices; we present results for G matrices estimated with the prior selected by comparison of the Deviance Information Criterion (DIC) scores (*V* = 0.5, *nu* = 2, see Table A2). We iterated models 10,100,000 times, and recorded values every 1000 generations with a burn-in period of 100,000 generations, for a total of 10,000 posterior estimates. We confirmed model convergence by examining trace and posterior distribution plots and calculated autocorrelations between samples (all of which were below the recommended level of 0.1) using the R package ‘coda’ (Plummer et al., 2006).

We calculated the mean genetic variances and genetic correlations from the posterior distribution of the 10,000 MCMC samples. Estimates of genetic correlations among standardized traits can range from -1 to 1, and we regarded genetic correlations as significant if their 95% highest posterior density intervals (HDPI) did not overlap 0. Estimates of genetic variance cannot be negative, so we determined the significance of these terms by comparing our estimates to those generated from null distributions where G matrices were fit to randomized data. First, we generated 1000 randomized data sets by randomly assigning phenotypes to individuals. We then fit G matrices to each of these randomized data sets and calculated the 95% confidence intervals of mean genetic variance for each of our four traits from these 1000 randomized G matrices. Estimates of genetic variance were considered significant if they exceeded the upper confidence limit of the null distribution. In addition to estimating genetic correlations among mutualistic traits, we calculated phenotypic correlation matrices as Pearson correlations among paired traits using the ‘psych’ package (Revelle, 2020), correcting for multiple tests in the determination of significance.

We calculated broad-sense trait heritabilities in the Jamaican metapopulation of *T. ulmifolia* by fitting linear mixed-effect models using restricted maximum likelihood (REML) in the ‘lme4’ package (Bates et al., 2015), and by implementing univariate Bayesian methods in MCMCglmm (Hadfield, 2010), using univariate versions of the animal models described above with modified priors (*nu* = 1). For the linear mixed models, we log-transformed total FN sugar mass but did not re-scale trait data. We modelled maternal line and block as random effects and we estimated broad-sense heritability as the proportion of total variance explained by maternal family. We used likelihood ratio tests implemented with the ‘RLRsim’ package (Scheipl et al., 2008) to test the statistical significance of our heritablity estimates by evaluating the significance of the random effect of maternal line.

Finally, to investigate the potential consequences of having selfed plants in the greenhouse for one generation on the expression of mutualistic traits, we measured out-crossed, selfed, and autogamous seed set in a subset of plants used to fit our G matrices. We incorporated selfed fitness data into a G matrix fit for the island metapopulation and regressed seed set against stigma-anther separation to investigate the possibility that our breeding design had a disproportionate effect on genotypes from out-crossing populations (see Appendix B for detailed methods).

#### Comparison of G matrices

To investigate differences in the patterns of genetic variance and covariance among individual populations of *T. ulmifolia*, we estimated G matrices for our five largest field populations (Brown’s Town, Cave, Mosquito Cove, Murdock, and Saint Ann’s Bay). We continued to standardize phenotype data to the grand mean and variance, rather than within each population separately, because this method maintains differences in trait means across populations (figure 3) that may be biologically meaningful (Hine et al., 2009; Puentes et al., 2016). We assessed potential differences in the overall size, or amount of total genetic variation among G matrices, by calculating the trace (the sum of diagonal elements) for each matrix and compared their 95% HPD intervals. We further explored variation in the orientation of G by calculating the eigenvectors and eigenvalues of our G matrices. Eigenvectors are linear vectors that capture the axes of greatest variation in multivariate space, and their corresponding eigenvalues reflect the relative amount of total variation they capture. By examining the eigenstructure of G, one can determine the combination of phenotypic traits that contributes most to multivariate variation in a dataset. To quantitatively test for differences among our G matrices, we used three methods: random skewers, Krzanowski’s common subspace analysis, and the fourth-order genetic covariance tensor (Hine et al., 2009; Aguirre et al., 2014; Chakrabarty and Schielzeth, 2020). Here, we discuss only the specifics of the fourth-order genetic covariance tensor, but detailed descriptions of the random skewers and Krzanowski’s common subspace analyses are in Appendix B.

**Figure 3:**
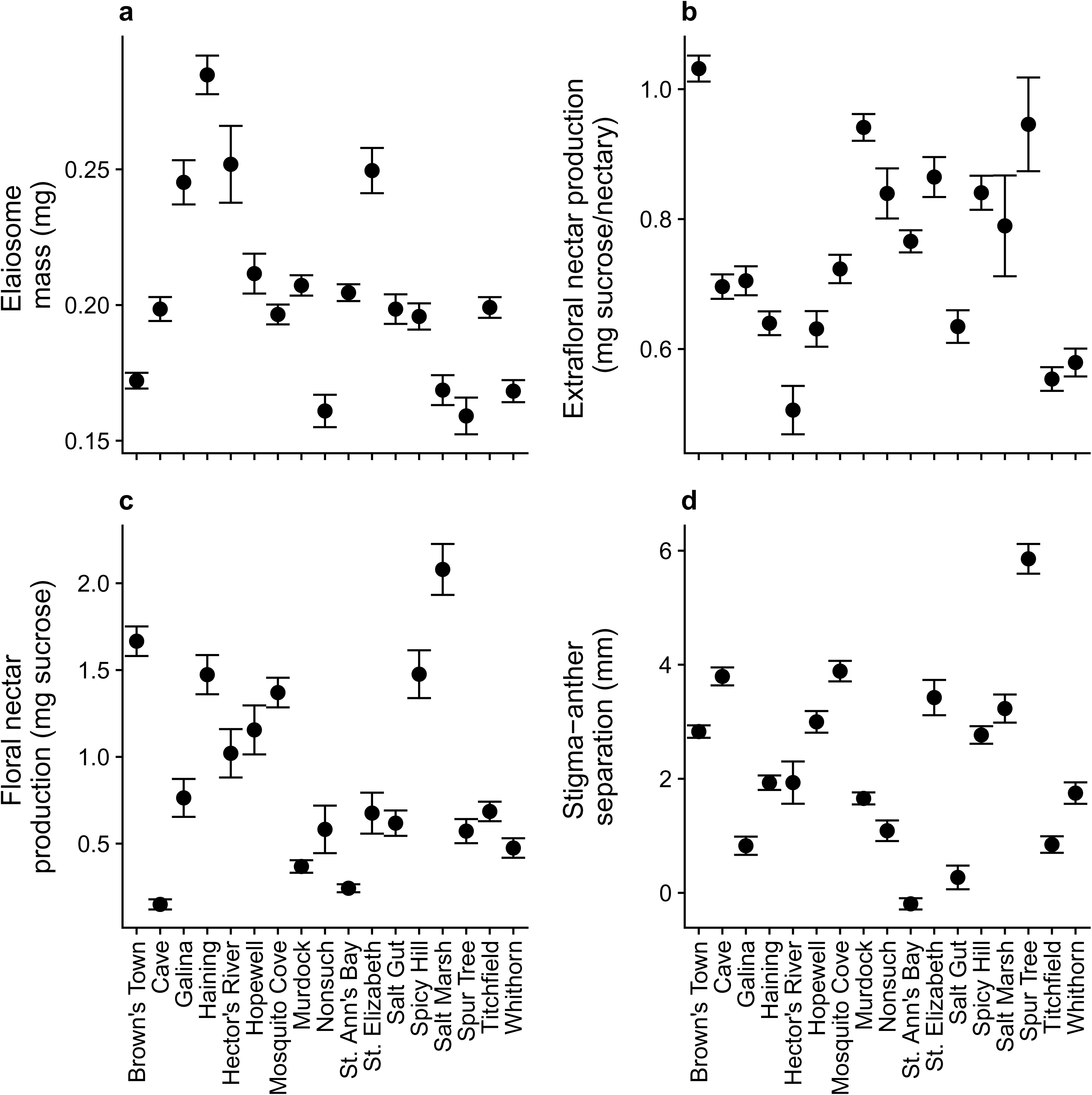
Variation in trait means (with standard error) across 17 *Turnera ulmifolia* populations.

The genetic covariance tensor method of Hine et al. (2009) allows for the comparison of G matrices directly by decomposing axes of variation among multiple matrices through eigenanalysis. Briefly, a tensor of the 4^th^ order is used to compare G matrices, which are themselves 2^nd^ order objects. This tensor is represented by Σ, and describes the variance and covariances among multiple covariance matrices. From this 4^th^-order covariance tensor, the second-order eigentensors, E that describe variation among matrices summarized by Σ can be explored and used to obtain the eigenvectors (e) and eigenvalues of E. The eigenvalues of an eigentensor can then be examined to explore which trait combinations contribute to detected changes in the covariance structures of individual G matrices. We employed the genetic covariance tensor method to test for and examine patterns of population divergence in G using code modified from Aguirre et al. (2014). We used Bayesian posterior estimates of G to project the eigentensors (E) onto Σ to determine 95% HPD intervals for *α*, the variance of Σ accounted for by each eigentensor. We then tested for significant differences among our underlying G matrices by comparing these estimates of *α* with those estimated from randomized G matrices. We generated randomized null G matrices for the purpose of significance testing by sampling and assigning breeding values (Aguirre et al., 2014) before reconstructing randomized individual phenotypes by incorporating empirical estimates of environmental variance. We then fit null G matrices using these phenotypes (Morrissey et al., 2019), and used the resultant randomized estimates of G to calculate null summary statistics for the genetic covariance tensor analysis and other G matrix comparison methods. Finally, we projected the leading eigenvectors (e) of the eigentensors (E) onto our observed G matrices in order to estimate the amount of genetic variance in the direction of the principal axes of multivariate genetic variation in each population, using all posterior estimates of G to fit 95% HPD intervals.

#### Evolutionary constraint and facilitation through G

To investigate the effects of genetic covariances on responses to selection in the island metapopulation and five largest populations of *T. ulmifolia*, we multiplied our estimates of G by three vectors of directional selection gradients representing simplified but plausible selection regimes. Measurements of selection on mutualistic traits in the field are rare, but our skewers reflect conservative estimates of selection based on published accounts. There are no estimates of selection on elaiosome mass, but their presence increases seed removal by ants by up to 20%, a large increase in fitness given that dispersal to ant nests can double germination rates in *T. ulmifolia* (Cuautle et al. 2005; Salazar-Rojas et al. 2012). Weak positive selection on EFN is also likely common, given that its presence boosts seed dispersal by a factor of up to 2.5 (Cuautle et al. 2005; Dutton et al. 2016b), and selection through plant defense can range from weakly negative (*β* = -0.096) to strongly positive (*β* = 0.695) depending on the presence of ants and herbivores (Rutter and Rausher 2004). Evidence of selection on floral traits is more mixed, as papers report little or no selection on FN volume and sugar content in flowering plants (e.g., Benitez-Vieyra et al. 2010; Kulbaba and Worley 2011; Gijbels *et al*. 2014), and positive selection on stigma-anther separation can be sex-specific (Kulbaba and Worley 2011). In all cases, we specified conservative skewers that featured either weak positive selection (*β* = 0.1), neutral selection (*β* = 0), or weak negative selection (*β* = -0.1) consistent with the range of published values.

In the first scenario, we specified universal, weak positive selection for all traits, while in the second selection regime, we simulated weak selection for defense (EFN) and dispersal (elaiosome mass), and against pollinator attraction (FN) and out-crossing (stigma-anther separation). In the third scenario, we simulated positive selection for defense and dispersal and against stigma-anther separation, but made FN production selectively neutral. We chose these three selection regimes because they represented ecologically plausible patterns of selection; namely, universal selection for investment in multiple mutualisms (1), selection for traits associated with dispersal, including concomitant selection for inducible defenses (EFN) and against floral traits associated with pollination and out-crossing (2), and selection for traits associated with selfing, with neutral selection on FN production given some evidence that it may be a relatively inexpensive resource (Harder and Barrett, 1992), and may not affect seed production in *T. ulmifolia* (Dutton et al., 2016a) (3).

We projected selection vectors through all 10,000 posterior estimates to produce response vectors, and calculated the 95% HPD intervals around the selection responses of individual traits for the purpose of significance testing. Additionally, we followed Agrawal and Stinchcombe (2009) to quantify the effect of genetic covariances on the rate of adaptation in our five bestsampled populations. Briefly, we compared the rate of adaptation with the observed G matrices to the rate of adaptation with the genetic covariances (off-diagonals) set to zero. In addition to calculating the responses to selection of our observed G matrices under our simulated selection scenarios, we calculated response vectors for their counterparts without genetic covariances. We then calculated R values by dividing the response vectors obtained with the full G matrix by those obtained with genetic covariances set to 0. R values greater than 1 indicate that genetic correlations facilitate adaptation, while values less than 1 indicate that adaptation is constrained by genetic correlations among traits (Agrawal and Stinchcombe, 2009).

In addition to assessing the impact of genetic correlations on the evolvability of mutualistic traits by measuring their effects on the response to selection, we also estimated the matrix of divergence among population trait means (D, Blows and Higgie 2003) and compared it to G_W_, the island-wide G. The eigenvectors or principal components of D represent the major multivariate axes of phenotypic variation across populations, and can be compared to the eigenvectors of G to determine whether or not the multivariate axes of genetic variation (e.g., genetic lines of least resistance, Schluter 1996) have constrained phenotypic divergence between populations. We estimated D by fitting a MCMCglmm model to the mean values of each of our four mutualistic traits for all 17 populations of *T. ulmifolia*, using the same number of MCMC samples and thinning intervals as those used in the fitting of G matrices. To compare G_W_ and D, we employed both one and two-dimensional comparison methods. We began by calculating the angle between the leading eigenvectors of each matrix (g_max_ and d_max_), to investigate how closely aligned the vectors describing maximal genetic and phenotypic differentiation are (Schluter, 1996; Blows and Higgie, 2003). Additionally, we explored the extent to which G has constrained phenotypic divergence among populations by comparing variation in the size and shape of G_W_ and D. For both summary matrices, we calculated the size of the matrices by summing their eigenvectors, and assessed their eccentricity by calculating the amount of variation captured by their leading eigenvectors. For each of these measures, we calculated the 95 % HPD intervals from the posterior estimates of our MCMCglmm objects. Finally, we employed the Krzanowski subspace analysis (described in Appendix B) to determine whether G_W_ and D share a common multivariate substructure (Silva et al., 2020).

## Results

### Genetic variances and covariances among traits

We found significant genetic variation for all four traits in the Jamaican metapopulation of *T. ulmifolia*; all genetic variance estimates in the metapopulation G matrix G_W_ (table 1), and all trait heritabilities calculated from the full data set using both REML and univariate Bayesian approaches (table A3) were statistically significant. The genetic variance in these traits was considerable, ranging from 0.17-0.77 depending on the trait and method of analysis (tables 1 and A3). All methods found that EFN production was the least heritable (0.17-0.22), albeit still with significant genetic variation among maternal lines, and stigma-anther separation was the most heritable trait (0.58-0.77), with FN production and elaiosome mass intermediate (0.48-0.61 and 0.46-0.55, respectively) (tables 1, A3, A4).

**Table 1:**
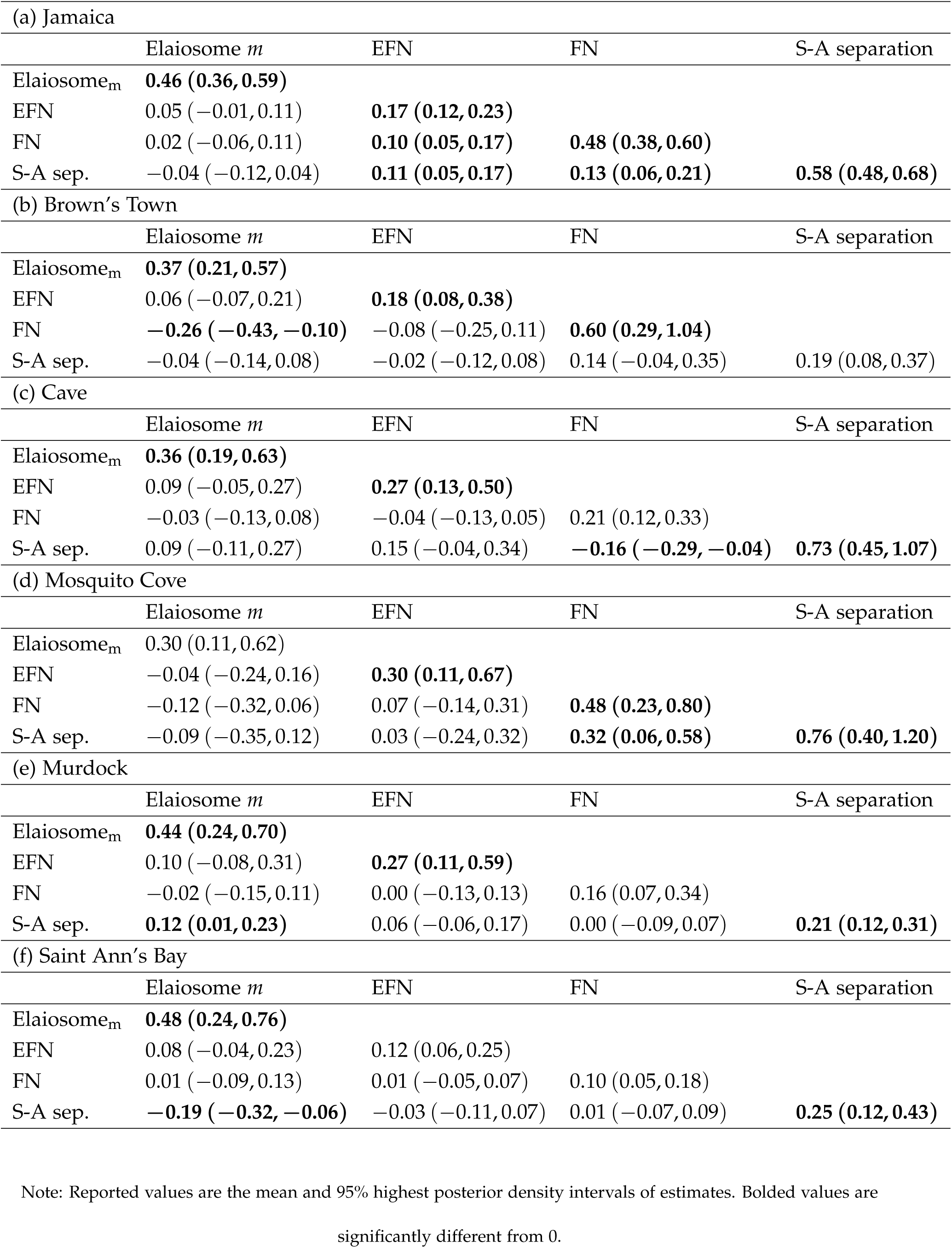
Genetic variance-covariance (G) matrices of the Jamaican metapopulation and five largest individual populations of *Turnera ulmifolia*

For the Jamaican metapopulation, all significant genetic correlations among traits were positive. We found that EFN and FN production were significantly genetically correlated, and both co-varied significantly with stigma-anther separation (table 1). In other words, maternal lines that secrete more EFN also make more herkogamous flowers that secrete more FN. In contrast, the genetic covariances between elaiosome mass and stigma-anther separataion, EFN, or FN production were small in magnitude (-0.04 to 0.05) and not significantly different from zero.

We found no evidence that selfing plants differentially affected genotypes from populations with higher stigma-anther separation. Although stigma-anther separation and seed set were negatively correlated across all treatments (table A6), we did not observe a significant interaction between pollen source and stigma-anther separation (figure A1, table A5). This analysis revealed a (positive) genetic correlation between selfed seed set and elaiosome mass, but no significant genetic correlations between seed set and the production of either EFN or FN, albeit in a analysis with low power for estimating G (table A6).

### Comparison of G matrices

Our fourth-order genetic covariance tensor analysis found no significant divergence among our five largest populations of *T. ulmifolia*. For each of the four eigentensors of the genetic covariance tensor, the 95% HPD intervals of *α* (the amount of genetic variance in the direction of each eigentensor) of the observed and randomized G matrices overlapped (figure 4A). Thus, these eigentensors do not describe significant variation among the observed G matrices of our five largest populations. These results are consistent with the results of both the application of random skewers and Krzanowski’s common subspace analysis, which both failed to detect significant differences among the G matrices of our largest populations (see Appendix A, figures A2, A3). Furthermore, the total amount of genetic variance in each of our population estimates of G did not differ significantly, as evidenced by their overlapping traces (see table 1). Taken together, these results suggest little divergence among our populations in the size and orientation of G.

**Figure 4:**
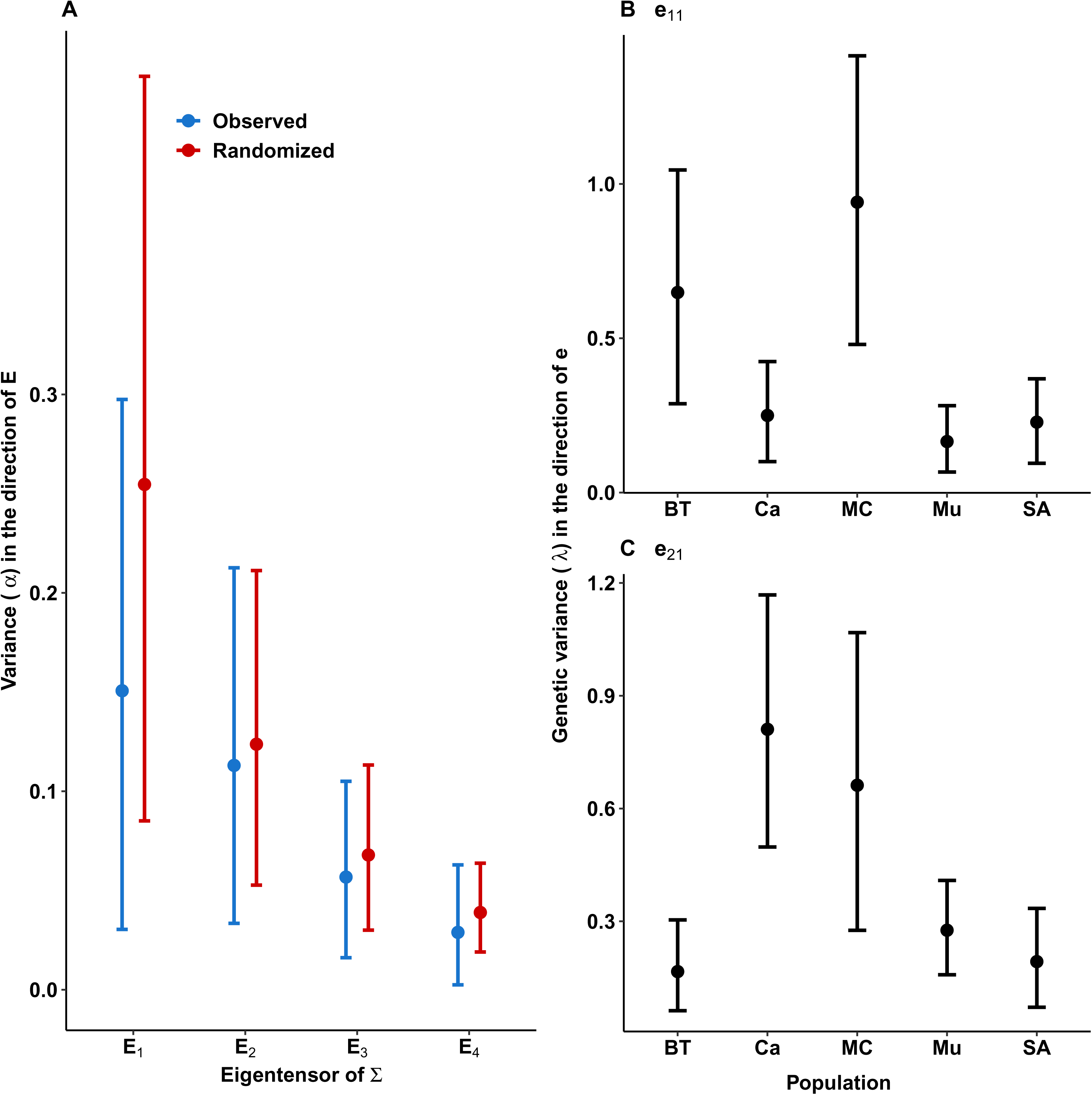
Results of the 4^th^ order genetic covariance tensor analysis comparing the G matrices of our five largest *Turnera ulmifolia* populations. (a) The amount of variance (*α*) in G explained by each eigentensor E of the 4^th^ order covariance tensor Σ for observed (blue) and randomized (red) G matrices. Genetic variance in the direction of (b) the first eigenvector of E1 and (c) the first eigenvector of E2. All error bars are 95% HPD intervals.

These comparisons, however, are frequently limited by issues of statistical power (Teplitsky et al., 2014; Puentes et al., 2016), particularly when G matrix estimates are fit with wide confidence intervals. Greater sampling may have shown differences in G among populations, given that projecting the leading eigenvectors of the first and second eigentensors of our covariance tensor (*e*_11_ and *e*_21_) back onto population-specific estimates of G revealed greater genetic variance in the Mosquito Cove and Brown’s Town populations (figure 4B) in the direction of *e*_11_ and in the Cave and Mosquito Cove populations in the direction of *e*_21_ (figure 4C) (see also Appendix B for more details). However, greater sampling in most *T. ulmifolia* populations was not possible, because of very small population sizes (table A1).

Our estimates of the phenotypic correlation (P) matrices for the individual populations and metapopulation of *T. ulmifolia* were broadly aligned with G. In all 36 paired trait combinations, the confidence intervals of our estimates of phenotypic and genetic correlations overlapped with one another (tables 1, A9). Because we grew plants and measured phenotypes in a common greenhouse environment, we expected that P and G matrices would be similar, and they were; we did not observe any cases where the sign of trait correlations were significant and different from one another.

### Evolutionary constraint and facilitation through G

Applying selection gradients onto the G matrices of the Jamaican metapopulation and our five largest populations revealed that the patterns of genetic correlation we observed can affect the adaptive potential of *T. ulmifolia*. When we simulated weak positive selection on all traits (scenario 1) in the Jamaican metapopulation, we found that positive correlations among extrafloral and floral nectar production and stigma-anther separation facilitated their response to simulated selection (mean R value with 95% HPD intervals: EFN, 2.51 (1.92-3.11), FN, 1.55 (1.27-1.83), stigma-anther separation, 1.36 (1.10-1.61), figures 5A, A4A). Unsurprisingly, these same correlations have the potential to dramatically reduce the visibility of mutualistic traits to multivariate selection when selection acts in opposing directions on floral traits and extrafloral nectar production. When we simulated a selection regime associated with selection for dispersal, inducible biotic defenses and increased selfing (2), we found that the evolution of extrafloral nectar was constrained (mean R value = 0.029, upper 95% HPD = 0.657, figure A4 B) to the point that it showed no response to positive selection (figure 5B).

**Figure 5:**
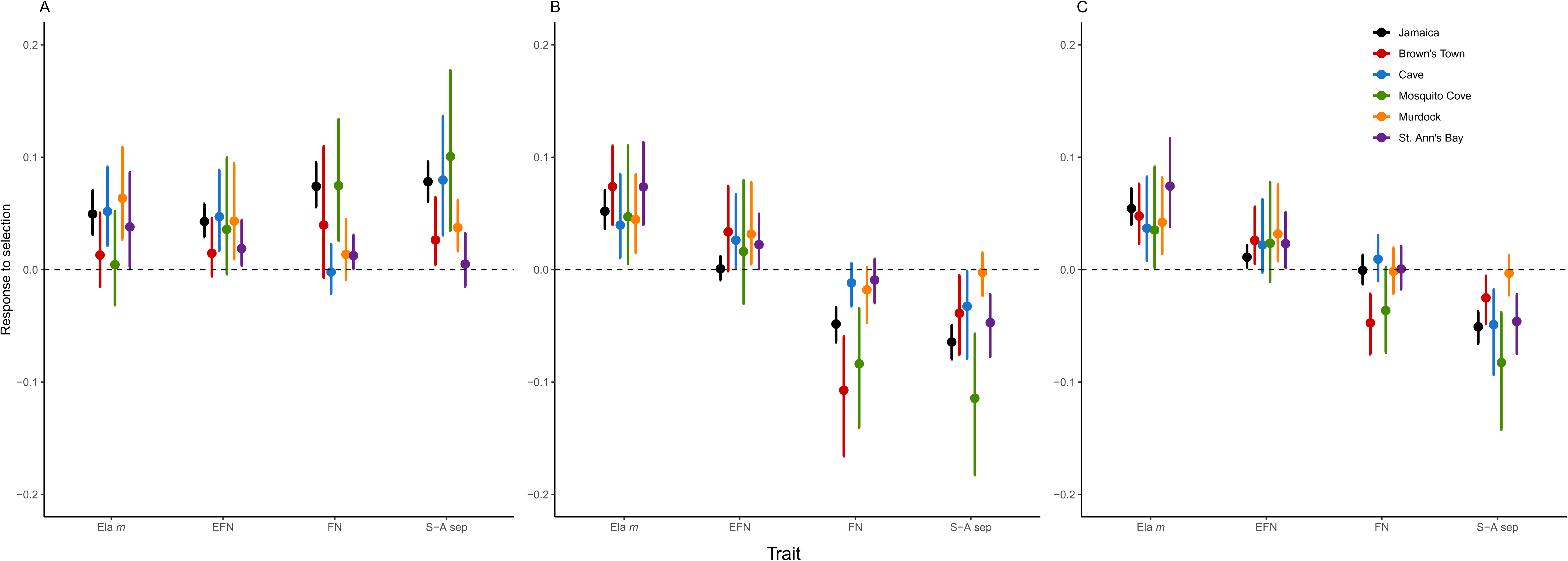
Responses of mutualistic traits to multivariate selection gradients in the Jamaican metapopulation and the five largest *Turnera ulmifolia* populations. We simulated selection scenarios in which (a) all traits are under positive directional selection (*β* = 0.1), (b) elaiosome mass (Ela *m*) and EFN are under positive directional selection while FN production and stigma-anther separation (S-A sep) are selected against (*β* = -0.1), and (c) elaiosome mass and EFN are under positive directional selection, S-A sep is selected against, and FN is selectively neutral (*β* = 0).

Among the five largest populations of *T. ulmifolia*, we found that slight but non-significant differences in G had the capacity to alter their responses to multivariate selection. In some cases, our analyses showed that population-specific estimates of G nearly completely inhibted their capacity to respond to positive directional selection, as was the case with floral nectar production in the Cave population and stigma-anther separation in Saint Ann’s Bay (figure 5A). In both cases, this result reflects a combination of low genetic variance for these traits coupled with significant negative genetic correlations with other traits under positive selection (table 1). Other populations, however, had more capacity to respond to selection on stigma-anther separation and floral nectar production due to the existence of positive genetic correlations that accelerated their response to selection (figure 5A). These results are consistent with estimated R values (see figure A4A), which provided evidence of genetic constraint in the evolution of FN production and stigma-anther separation in the Cave and Saint Ann’s Bay populations. We observed similar variation in the capacity of our five populations to respond to multivariate selection that exerted opposing pressures on mutualistic traits. Here again, we found that misalignment between axes of genetic correlation and selection vectors reduced the capacity for trait evolution in certain populations (e.g., floral nectar in Cave, Murdock and Saint Ann’s Bay), while the responses to selection for this same trait in other populations (Brown’s Town and Mosquito Cove) were facilitated by the existence of either negative correlations among traits experiencing conflicting selection pressures or positive correlations among traits under the same selection pressures (figure 5B,C, table 1). We observed similar variation in the effects of genetic correlations on the evolution of stigma-anther separation, where alignment between axes of selection and co-variation facilitated its response to selection in the Brown’s Town, Mosquito Cove, and Saint Ann’s Bay populations and misalignment constrained its evolution in the Murdock population (figure 5 B,C, A4).

When we compared the size, structure, and orientation of G_W_ and D, the matrix summarizing multivariate divergence among the mutualistic traits of all 17 *T. ulmifolia* populations, we found that the structure of G_W_ has constrained phenotypic divergence at the metapopulation level. There was little difference in the size of the D (mean with 95% HPD intervals: 1.91 (1.12-2.82)) and G_W_ (1.69 (1.48-1.88)) matrices, and also little difference in their eccentricities (0.408 (0.374-0.674) and 0.422 (0.379-0.489) for D and G_W_, respectively). The multivariate directions of maximal genetic (g_max_) and phenotypic (d_max_) variation were also very closely aligned, with an angle of only 2.80°(0.219°-2.91°) between them. When we assessed the extent to which our estimates of G_W_ and D shared a common substructure using Krzanowski’s shared subspace analysis, we found further evidence of similarity. The eigenvalues of the eigenvectors of H, the summary matrix encapsulating the shared multivariate subspace between matrices, overlapped with the maximal value of 2 (*p*, the number of matrices being compared) for both interpretable eigenvectors (h_1_: 1.85 (1.49-2.00), h_2_: 1.60 (0.99-2.00)). This result suggests a shared subspace among our G_W_ and D matrices (Krzanowski, 1979; Aguirre et al., 2014). Thus, G_W_ and D do not differ in size, eccentricity or orientation.

## Discussion

Genetic correlations among mutualist reward traits in *T. ulmifolia* have the capacity to affect the evolution of mating system and rewards for both pollinators and biotic defenders. We found significant positive genetic correlations among the production of FN, EFN, and stigma-anther separation in the common genetic variance-covariance matrix G_W_ for the Jamaican metapopulation of *T. ulmifolia*. In other words, floral nectar production was coupled to extrafloral nectar production, and both were linked to variation in plant mating system, while elaiosome mass was not genetically correlated with other mutualistic traits. At the island-wide level, these positive genetic correlations are predicted to accelerate *T. ulmifolia*’s response to selection when selection favours increased investment in pollination, out-crossing, and biotic defense. However, opposing selection pressures on floral traits and biotic defense can constrain the evolution of mutualistic reward traits in *T. ulmifolia*. The multivariate axes of phenotypic divergence among our *T. ulmifolia* populations were closely aligned with the prominent axes of multivariate genetic variation in the island metapopulation, suggesting that the structure of G has constrained phenotypic evolution in the *T. ulmifolia* metapopulation. However, there were few differences in the orientation of G among populations.

### Genetic variances and covariances among mutualistic traits

Selection on traits associated with multiple mutualisms likely varies across space and time given that ecological outcomes are affected by community composition and non-additive interactions among partners affect fitness outcomes of their hosts (Afkhami et al., 2014). The outcome of multiple mutualisms involving pollinators, biotic defenders, and seed-dispersing ants may be particularly prone to non-additive interactions that alter the pattern of multivariate selection. Floral and extrafloral nectar are biochemically or physiologically linked (Heil, 2011; Dutton et al., 2016a), and selection on floral traits associated with out-crossing and pollination can depend on both the presence of ant bodyguards that deter pollinators (Cembrowski et al., 2014; Villamil et al., 2019) and herbivores (Knauer and Schiestl, 2017; Ramos and Schiestl, 2019) that are in turn affected by the presence of ants. As such, our results may help explain the ubiquity and stable co-existence of both pollination by insects (Ollerton et al., 2011) and ant defense (Marazzi et al., 2013; Weber and Keeler, 2013), despite physiological trade-offs and negative ecological interactions among partner species. A positive genetic correlation between floral and extrafloral nectar production may serve to maintain both mutualisms by reducing their visibility to negative selection and indirectly leading to the evolution of increased reward production through selection on correlated traits, even when the benefits of a particular mutualism may be context-dependent or transient.

Our findings also hint at potential mechanisms responsible for generating genetic correlations between floral and extrafloral nectar production. Given the biochemical similarity of floral and extrafloral nectar (Heil, 2011), a positive correlation suggests shared genes are involved in nectar synthesis. Alternatively, a positive genetic correlation between floral and extrafloral nectar production might itself be adaptive. While extrafloral nectaries are thought to serve several adaptive functions, namely the attraction of bodyguards that protect against herbivory and, in our system, the attraction of seed-dispersing ants (Janzen, 1966; Cuautle and Rico-Gray, 2003; Cuautle et al., 2005; Heil, 2011; Dutton et al., 2016b), the distraction hypothesis has long been proposed as an alternate explanation for the evolution of extrafloral nectaries (Kerner, 1878). The distraction hypothesis posits that extrafloral nectar production is adaptive because it incentivizes ant recruitment away from flowers, thereby reducing ant consumption of floral nectar and ant-pollinator conflict (Cembrowski et al., 2014; Villamil et al., 2018, 2019). Hence, a positive genetic correlation between floral and extrafloral nectar production in *T. ulmifolia* could be the result of pleiotropy, or of strong selection on plants with showy and nectar-rich flowers to prevent ants from interfering with the pollination process.

Similarly, there may be an adaptive explanation for the positive genetic correlations we observed between stigma-anther separation and both floral and extrafloral nectar production in the Jamaican metapopulation of *T. ulmifolia*. Plants face a trade-off between attracting pollinators and floral antagonists with showy floral displays, and multiple studies have shown that variation in floral morphology and mating system is more visible to selection when herbivores are absent (Thompson and Johnson, 2016; Knauer and Schiestl, 2017; Ramos and Schiestl, 2019). Both the deterrence of herbivores and the isolation of ant bodyguards away from floral tissues may thus be of greater selective value to genotypes with high stigma-anther separation, a quantitative trait correlated with out-crossing rate in *T. ulmifolia* (Belaoussoff and Shore, 1995). Correlated selection on suites of traits such as this can lead to the evolution of stronger genetic correlations among them (Jones et al., 2004, 2012; Penna et al., 2017). Positive genetic correlations between stigma-anther separation, and pollinator and biotic defender rewards in *T. ulmifolia* are also consistent with the positive correlations between floral traits and extrafloral nectar production across species of *Gossypium* (Chamberlain and Rudgers, 2012). Such positive correlations may reflect a broader pattern of correlated selection on mating-system and biotic defense in flowering plants.

### Comparison of G matrices

As with other genetic entities, variation in G can arise through familiar evolutionary forces including heterogeneous selection, drift, migration, and mutation (Roff, 2000; Jones et al., 2004; Guillaume and Whitlock, 2007; Bjö rklund and Gustafsson, 2015; Hangartner et al., 2020). That we did not observe divergence among the G matrices of our five largest populations of *T. ulmifolia* suggests that the structure of G is relatively stable to these perturbations across Jamaica. These results are consistent with those of other studies that have reported stability in the shape and size of G within a single species, even across large geographic and climatic ranges (e.g., Teplitsky et al., 2011; Puentes et al., 2016; Sniegula et al., 2018; Hangartner et al., 2020).

We must, however, interpret evidence of similarity in G with some caution given the relatively low number of individuals (153–167) and maternal families (19–20) used to fit the G matrices of our largest populations and the conservative nature of G matrix comparison methods (Puentes et al., 2016; Sniegula et al., 2018). Local populations of *T. ulmifolia*, a highly opportunistic weed, tend to be small and ephemeral (table A1, Barrett, 1978; Barrett and Shore, 1987); in many cases, we collected seed from all or nearly all individuals in a natural population of *T. ulmifolia*, meaning we could not have phenotyped plants from additional maternal families for these populations. Although all methods found no significant differences in G among populations, our populations nonetheless differed in their amount of genetic variance along the principal axes of multivariate genetic variation captured by the fourth-order covariance tensor (figure 4B and C), and there were meaningful differences in evolutionary responses to selection among some populations (e.g., FN production in Cave and Mosquito Cove, figures 5A and A4). The movement of *T. ulmifolia* seeds across the landscape is facilitated by secondary seed dispersal by ants and its use as an ornamental by humans (Gómez and Espadaler, 1998; Bottcher et al., 2016). Bottleneck events such as those associated with stochastic colonization events can affect the structure of G (Whitlock et al., 2002), and affect the relative genetic contribution of local populations to a connected meta-population (e.g., hard selection; Wade, 1985; De Lisle and Svensson, 2017). However, local variation in G may not persist in the face of extensive gene flow in a connected metapopulation.

### Evolutionary constraint and facilitation through G

When ecologically relevant traits are genetically correlated, as in *T. ulmifolia*, the evolutionary responses of populations are determined not only by genetic variance in and selection on individual phenotypes, but also by the genetic covariances among traits, which can facilitate or constrain adaptive evolution depending on whether axes of genetic covariance are aligned with those of selection (Agrawal and Stinchcombe, 2009; Wise and Rausher, 2013; Teplitsky et al., 2014; TerHorst et al., 2018). We found that the evolution of FN, EFN, and stigma-anther separation will be facilitated when selection favours increased investment in pollination, out-crossing, and biotic defense (figures 5 A, A4A). Conversely, under selection regimes consistent with selection for selfing, favouring increased inducible defenses and dispersal and reduced FN production and stigma-anther separation (Baker, 1955; Belaoussoff and Shore, 1995; Pannell and Barrett, 1998; Campbell and Kessler, 2013; Campbell et al., 2014), we found that both EFN and FN showed little or no response to simulated selection in the Jamaican metapopulation of *T. ulmifolia* (figure 5B and C, figure A4B and C). In other words, plant mutualisms with pollinators and ants are unlikely to evolve independently in *T. ulmifolia*, but whether adaptive evolution is accelerated or hindered by G will depend on the nature of multivariate selection. The response to directional selection acting on one trait would, for example, differ depending on whether stabilizing selection is resisting change in other correlated traits (‘i.e., conditional evolvability,’ Hansen and Houle, 2008).

Although G did not differ significantly among populations, there was extensive variation in the means of multiple mutualism-associated traits among populations (figure 3), consistent with a pattern of genetic constraint on the direction of phenotypic evolution. The matrix of multi-variate phenotypic divergence among population trait means, D, and G_W_ were similar in size, eccentricity, and orientation, and there was only a 2.80° angle between the directions of maximal genetic variation (g_max_) and phenotypic divergence (d_max_). These results may be evidence of genetic constraint, with phenotypic divergence among populations being restricted to genetic lines of least resistance, i.e., the prominent multivariate axes of genetic variation (Schluter, 1996). Gene flow across the island of Jamaica may also explain the alignment we observed between patterns of multivariate genetic variance and the direction of phenotypic divergence; simulations have shown that high gene flow can realign G by increasing its genetic variance in the direction of phenotypic divergence (Guillaume and Whitlock, 2007). Multiple previous studies have demonstrated that G constrains phenotypic divergence among populations, including in populations separated by greater evolutionary (e.g., McGlothlin *et al*., 2018) and geographic (e.g., Silva et al., 2020) distances than ours.

### Conclusion

The evolution of traits mediating multiple species interactions will be influenced by both ecological complexity and their underlying genetic architectures. Positive genetic correlations among FN, EFN, and stigma-anther separation in *T. ulmifolia* have the potential to link the evolutionary fates of these traits in a metapopulation. These findings highlight the importance of estimating the genetic correlations among traits characterized by ecological (Cembrowski et al., 2014; Villamil et al., 2018; Ramos and Schiestl, 2019; Villamil et al., 2019) and physiological trade-offs (Dutton et al., 2016a). Despite the potential for conflict between pollination and biotic defense, we found that the production of nectar rewards for these two mutualisms was positively correlated at the metapopulation level, and uncorrelated within individual smaller populations. Genetic correlations among mutualistic traits may be a result of pleiotropy, but may also plausibly reflect a process of coordinated positive selection on a suite of phenotypes. These patterns appear to be relatively stable in Jamaican *T. ulmifolia*, as we found weak evidence of local variation in G among populations across the island. The stability of G suggests that local variation is unlikely to impact the evolution of multiple mutualisms at the metapopulation level, and that phenotypic divergence in mutualistic traits may be constrained by the prominent axes of multivariate genetic variation in a metapopulation. Our results provide rare insight into the genetic architecture of multiple mutualisms, highlighting the potential for genetic correlations among reward traits to affect the evolution and ecology of these commonplace and essential interaction networks.

## Acknowledgements

We would like to thank many undergraduate students for their assistance in plant care and data collection, including A. Copeland, A. Wichert, B. Garden-Smith, C. Chen, C. Qiao, E. Zhang, F. Samada-zada, J. Han, K. Ong, M. Giblon, M. Gorchkova, S. Fernandes, and S. Ravoth; S. Barrett for guidance in planning field collections; J. Shore and E. Dutton for advice on working with *Turnera ulmifolia*; J. Stinchcombe, J. Sztepanacz, and C. Wood for assistance in fitting and comparing G matrices; J. Stinchcombe for providing comments on our manuscript; and A. O’Brien and B. McGoey for helping with coding. M.E.F. received funding from an NSERC Discovery Grant and the University of Toronto; J.R.L. was supported by an NSERC CGS-D Alexander Graham Bell scholarship; C.G.R. was supported by an NSERC Undergraduate Student Research Award. The authors declare no conflicts of interest.

## Author contributions

JRL and MEF designed the study. JRL collected seeds and grew plants; JRL, CGR, CB, TW, and CK collected trait data. CGR, JRL, and MEF designed the hand-pollination experiment; CGR collected data. JRL analysed data and drafted the manuscript; JRL and MEF finalized the text.

## Appendix A: Supplementary Materials and Methods

### Calculation of nectar concentration and sugar content

We calculated the concentration of the original extrafloral (EFN) and floral (FN) nectars of plants using the following equation:

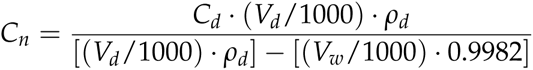

where *C_n_* and *C_d_* are the concentrations (g sucrose/ 100 g solution) of the nectar and the diluted samples, respectively, *V_d_* is the volume of the diluted sample (*µ*L), *ρ_d_* is the density of the diluted sample (g/mL), *V_w_* is the volume of water we added to dilute the sample (*mu*L), and 0.9982 is the density of distilled water (g/mL). We used a sucrose density chart to estimate the density of our sample (Haynes and Lide, 2010).

We estimated the total sugar mass of EFN and FN samples using the following equation:

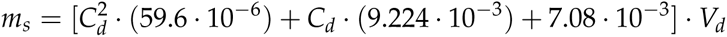

where *m_s_* is sucrose mass (mg) of the nectar sample, *C_d_* is the concentration of the diluted sample (g sucrose/ 100 g solution), and *V_d_* is the volume of the diluted sample (*µ*L). This is a quadratic equation of the relationship between nectar concentration by weight and by volume (Bolten et al., 1979; Burquez and Corbet, 1991; Dutton et al., 2016a). For nectar concentrations that range between 10% and 80%, as in nearly all of our diluted EFN and FN samples, the error produced by this equation is expected to be less than 1% (Burquez and Corbet, 1991; Dutton et al., 2016a).

### Fitness consequences of mating system variation

We hand-pollinated a subset of plants to investigate the fitness consequences of putative variation in mating system. After collecting trait data, we haphazardly chose three plants from each of five randomly chosen maternal lines per population, for a total of 255 plants from 85 maternal lines.

We applied three hand-pollination treatments to each plant: (1) out-crossed, where flowers were hand-pollinated using a random pollen donor from the same population, but different maternal line; (2) selfed, where flowers were hand-pollinated by applying pollen from its own anthers; and (3) autogamous, where flowers were not manipulated and pollen transfer occurred upon withering of the perianth (Barrett, 1978). We measured stigma-anther separation for each experimental flower in all three treatments, and emasculated selfed and out-crossed flowers to prevent unintended pollen transfer. We did not observe any potential pollinators in the greenhouse over the course of this experiment. Twenty-one days after we pollinated flowers, we collected mature fruits and counted total seed production.

To investigate the effects of pollination treatment and variation in mating system on seedset, we used the lme4 package (Bates et al., 2015) to fit a linear mixed effect model. We log-transformed seed set data and modelled the interaction between stigma-anther separation and pollination treatment as fixed effects. We included random effects for individual plant nested within maternal line and population, as well as for the interaction between population and pollination treatment. We assessed the significance of this population by treatment interaction using likelihood ratio tests in the RLRsim package (Scheipl et al., 2008), and performed post-hoc tests using the emmeans package (Lenth, 2019) to compare the effects of different pollination methods on seed set.

Furthermore, we investigated the consequences of our breeding design by fitting a G matrix that included our original four phenotypes of interest (elaiosome mass, EFN, FN and stigma-anther separation) along with selfed seedset.

### Random skewers

The random skewer method assesses whether G matrices diverge from one another by determining whether their evolutionary responses to randomly generated selection vectors differ (Roff et al., 2012). We compared the evolutionary responses of each of our 10,000 posterior estimates of G for our 5 largest *T. ulmifolia* populations to 10,000 randomly generated selection vectors following the methods of Roff and colleagues (2012). The selection vectors consisted of four selection vector elements drawn randomly from a normal distribution centered at 0 with a standard deviation of 1 and standardized to generate a continuous distribution of negative and positive selection vectors biased towards weak to moderate directional selection (Aguirre et al., 2014). For each vector, we calculated the multivariate response to selection for each posterior MCMC estimate of G for each population. We then calculated response vector correlations for each pair of populations at each posterior estimate of G in order to estimate mean response vector correlation for each population pair for each vector. We used these 10,000 estimates of mean response vector correlations to calculate the 95% HPD intervals in order to compare the similarity of evolutionary responses to random selection vectors among our populations. To determine whether or not the response vector correlations among populations differed from the null hypothesis of no difference in G, we compared the 95% HPD intervals of the response vector correlations of our G matrices against those estimated from null G matrices fit on phenotypes reconstructed from randomized breeding values and environmental variance (Morrissey et al., 2019).

We also implemented a modified version of random skewers developed by Aguirre and colleagues (2014), designed to estimate the amount of genetic variance in the direction of each random skewer in G. Briefly, we projected 10,000 random vectors through all posterior estimates of G for our 5 populations in order to estimate a posterior distribution of genetic variance in the direction of each of these vectors. We then deemed populations to differ significantly from one another in their genetic variances in the direction of a particular random skewer if the HPD intervals of their genetic variance did not overlap. Random vectors that produced significant variation among populations were then be collated, and the product-moment of their vector components calculated to create a 4 x 4 matrix, R, that encapsulates the elements of phenotypic space that show evidence of significant differences in genetic variance among populations. We then evaluated the most important axes of variation of R through eigenanalysis and projected its eigenvectors back onto the G matrices of our five populations to estimate the amount of genetic variance in the direction of each of these eigenvectors (Aguirre et al., 2014). We deemed populations as significantly different in their patterns of genetic variance in the direction of these eigenvectors if their 95% HPD intervals of genetic variance did not overlap.

### Krzanowski’s common subspace analysis

Krzanowski’s common subspace analysis can be used to determine whether the genetic sub-spaces of G that contain the most genetic variation are shared among multiple populations (Krzanowski, 1979). This method, precisely because it restricts itself to investigating the sub-space of G with the most variation (ie. the subspaces that will most influence responses to selection) speaks not only to shared genetic subspaces among populations, but reflects how differently populations can be expected to evolve under the same selection pressures (Aguirre et al., 2014). We assessed the extent to which our populations shared a common genetic subspace that encompasses most of their genetic variation by following and modifying code from Aguirre et al. (2014). Briefly, we calculated a summary matrix H containing the eigenvalues of a subset *k* of the eigenvectors that explain most of the shared genetic variation among populations. Eigen-values of H can take on a maximum value of *p*, the number of matrices being compared, with values equivalent to *p* indicating that these eigenvalues describe genetic variation shared among all populations. In contrast, eigenvalues less than *p* reflect divergence of at least one population among those sampled. For eigenvalues that describe axes of genetic variation not shared among all populations, one can quantify the angle between the genetic subspace of populations and the relevant eigenvector of H (Aguirre et al., 2014). We assessed the common subspace of G based on the standard specification of *k*, the subset of eigenvectors of each population’s G matrix used to construct H (*k* half the number of traits). To test for the significance of eigenvectors explaining differences in genetic variation among populations, we assessed whether the 95% HPD intervals of the eigenvalues of H overlapped with *p*, and compared these results to those from randomized G matrices fit on simulated phenotypic data reconstructed from randomized breeding values and environmental variance (Morrissey et al., 2019).

## Appendix B: Supplementary Results and Figures

### Fitness consequences of mating system variation

Across all three pollination treatments, stigma-anther separation had a significant negative effect (Type III Wald test, *χ*^2^ = 7.22, *P <*0.01, Table A5) on seed production (figure A1). Pollen source significantly affected mean seed set (Type III Wald test, *χ*^2^ = 7.22, *P <*0.0001) and flowers in the autogamous treatment (pollination through wilting of the perianth) produced significantly fewer seeds than flowers to which we applied pollen directly. However, whether we supplied flowers with pollen from their own anthers (selfed) or from flowers belonging to a plant from the same population (out-crossed) had no effect on seed set and the effect of stigma-anther separation on seed set was consistent across all treatments (Table A5). While the interaction between stigma-anther separation and pollination treatment was non-significant, the effects of pollination treatment on seed set did vary among different *T. ulmifolia* populations (RLRT = 17.02, *P <*0.0001, Table A5).

Variation in autogamous seed production was not genetically correlated with stigma-anther separation, EFN or FN production, although the dataset is quite small for estimating a G-matrix. We did find a significant positive genetic correlation between selfed seed set and elaiosome mass (Table A6).

### Random skewers

The response vector correlations among our five largest *Turnera ulmifolia* were moderately low, ranging from mean values from 58.2% to 76.6%. The 95% HPD intervals of all 10 correlation estimates overlapped with those estimated on randomized G matrices, suggesting the absence of significant differences in G between all population pairs.

When we implemented a modified approach to random skewers that estimates the amount of genetic variance in the direction of each vector (Aguirre et al., 2014), we found that 4,424 of 10,000 random vectors resulted in significant differences between two or more populations. We then calculated the product-moment correlation of the 4,424 vectors that captured significant differences in G among our populations to generate a matrix, R, that represents the area of phenotypic space characterized by significant differences in genetic variance among populations. The two leading eigenvectors of R, *r*_1_ and *r*_2_, accounted for 40.6% and 35.8% of the total variation, respectively. Only these two eigenvectors resulted in HPD intervals of genetic variance that did not overlap between at least two populations (figure A2). The first eigenvector, *r*_1_ reflects a trait combination dominated by variation in stigma-anther separation, and that displays much greater genetic variance in the Cave population and significantly less genetic variance in the Brown’s Town, Murdock, and Saint Ann’s Bay populations. The second eigenvector, r_2_ corresponded to a combination of traits primarily driven by floral nectar production, and showed higher genetic variance in Mosquito Cove and Brown’s Town, than it did in the Cave, Murdock, and Saint Ann’s Bay (figure A2). The results we obtained from the modified application of random skewers were very similar to those from the eigenanalysis of the 4th-order genetic covariance tensor. The amount of genetic variance in the direction of the leading eigenvector of the first eigentensor of Σ, *e*_11_ was very similar to the second eigenvector of R, *r*_2_. Likewise the amount of genetic variance in the direction of *e*_21_ was remarkably consistent with the first eigenvector of R, *r*_1_ (figure 4, A2).

### Krzanowski’s common subspace analysis

The results of Krzanowski’s common subspace analysis yielded similar results to those we obtained by applying random skewers and the fourth-order genetic covariance tensor to the G matrices of our five largest populations of *Turnera ulmifolia*. The first two eigenvectors of each G explained only 70.7–78.9% of the genetic variance observed in our G matrices, a relatively modest amount. The eigenvalues of these eigenvectors were both significantly smaller than the maximal value of *p* (5) (figure A3). This indicates that for at least one population, the eigenvalues of these eigenvectors of H (the genetic subspace shared among the populations) cannot be reconstructed from the eigenvectors of its specific G matrix (Krzanowski, 1979; Aguirre et al., 2014). This, however, cannot be interpreted as robust evidence of divergence among G matrices, and when we compared the eigenvalues of our observed G matrices to those extracted from comparisons of our randomized G matrices, the 95% HPD intervals overlapped in all cases (figure A3). Thus, these results are consistent with those obtained from other methods of G matrix comparison, and indicate that differences in the G matrices of our 5 largest populations of *T. ulmifolia* are limited.

### Additional results from fourth-order genetic covariance tensor analysis

When we estimated the genetic variance of each population along the prominent multivariate axes of variation extracted from summary matrices encapsulating variation among populations, we found that their 95% HPD intervals did not always overlap. For the leading eigenvectors of both the first and second eigentensors of our covariance tensor (*e*_11_ and *e*_21_), we found that populations differed in the amount of genetic variance aligned with these multivariate axes of variation. The leading eigenvector of the first eigentensor, *e*_11_, captures multivariate variation dominated by floral nectar production and stigma-anther separation. When we projected it back onto population-specific estimates of G, it revealed greater genetic variance in the Mosquito Cove and Brown’s Town populations (figure 4B). Likewise, the projection of *e*_21_, which reflects variation in stigma-anther separation and extrafloral nectar production, demonstrated greater genetic variance in the Cave and Mosquito Cove populations along this axis (figure 4 C). These results are strikingly similar to the projection of the leading eigenvectors of the summary matrix, R, that reflects the phenotypic space of greatest genetic variance as determined by the application of random skewers (see figure A2). While these results should be interpreted with caution given the lack of significant differences in G, they suggest a trend towards differentiation whose detection is limited by low statistical power in our analyses.

**Table A1:** Locations and sizes of *Turnera ulmifolia* populations

| Population | Longitude | Latitude | Population size | Maternal lines | Individuals |
| --- | --- | --- | --- | --- | --- |
| Haining | 17.99 N | 76.28 W | 22 | 12 | 111 |
| Nonsuch | 18.15 N | 76.38 W | 4 | 4 | 33 |
| Titchfield | 18.18 N | 76.45 W | 15 | 15 | 120 |
| Hector's River | 18.01 N | 76.27 W | 4 | 2 | 19 |
| Salt Gut | 18.41 N | 76.99 W | 7 | 6 | 51 |
| Murdock | 18.41 N | 76.92 W | 38 | 19 | 156 |
| Galina | 18.40 N | 76.89 W | 7 | 7 | 61 |
| Spur Tree | 18.01 N | 77.58 W | 8 | 8 | 63 |
| Saint Elizabeth | 18.08 N | 77.75 W | 7 | 7 | 62 |
| Cave | 18.21 N | 78.05 W | 35 | 20 | 162 |
| Mosquito Cove | 18.46 N | 78.14 W | 41 | 20 | 153 |
| Hopewell | 18.45 N | 78.03 W | 7 | 7 | 63 |
| Whithorn | 18.26 N | 78.03 W | 14 | 13 | 95 |
| Salt Marsh | 18.50 N | 77.71 W | 17 | 10 | 40 |
| Saint Ann's Bay | 18.43 N | 77.19 W | 38 | 19 | 162 |
| Brown's Town | 18.39 N | 77.36 W | 26 | 20 | 167 |
| Spicy Hill | 18.45 N | 77.55 W | 8 | 8 | 72 |

Note: We report the number of individual plants fully phenotyped in the greenhouse, and the number of maternal lines from each site from which individuals were phenotyped.

**Table A2:** Specification and DIC scores of G matrices estimated with different priors

| $V$ | $nu$ | DIC score |
| --- | --- | --- |
| 0.5 | 4 | 11514.35 |
| 0.333 | 4 | 11664.03 |
| 0.25 | 4 | 11723.61 |
| 0.5 | 2 | 11260.37 |
| 0.5 | 3 | 11414.48 |
| 0.25 | 2 | 11471.34 |
| 0.25 | 3 | 11621.60 |
| 0.333 | 2 | 11411.38 |
| 0.333 | 3 | 11411.38 |

Note: Inverse Wishart priors varied in expected genetic variance $V$ and degree of belief, $nu$ .

**Table A3:** Trait-specific broad-sense heritabilities for the Jamaican metapopulation of *Turnera ulmifolia*

| Estimator | Elaiosome $m$ | EFN | FN | S-A separation |
| --- | --- | --- | --- | --- |
| REML | **0.442** | **0.178** | **0.478** | **0.606** |
| Bayesian | **0.550** | **0.216** | **0.611** | **0.774** |

Note: We calculated heritabilities using restricted maximum likelihood (REML) and Bayesian methods. Bolded values are statistically significant at $p < 0.05$ .

**Table A4:** Linear mixed model results partitioning variance among the random effects for *Turnera ulmifolia* traits in the island-wide metapopulation

| | Elaiosome $m$ | EFN | FN | S-A separation |
| --- | --- | --- | --- | --- |
| Unnested model: |  |  |  |  |
| Maternal line | $1.43 \cdot 10^{-3}$ | $3.00 \cdot 10^{-3}$ | $9.35 \cdot 10^{-2}$ | 2.91 |
| Block | $5.19 \cdot 10^{-6}$ | $1.01 \cdot 10^{-4}$ | $4.68 \cdot 10^{-4}$ | $3.16 \cdot 10^{-2}$ |
| Residual | $1.80 \cdot 10^{-3}$ | $1.49 \cdot 10^{-2}$ | $1.02 \cdot 10^{-1}$ | 1.86 |
| Nested model: |  |  |  |  |
| Maternal line : Population | $5.36 \cdot 10^{-4}$ | $1.50 \cdot 10^{-3}$ | $1.16 \cdot 10^{-2}$ | $6.09 \cdot 10^{-1}$ |
| Population | $1.13 \cdot 10^{-3}$ | $1.48 \cdot 10^{-3}$ | $7.52 \cdot 10^{-2}$ | 2.37 |
| Block | $5.03 \cdot 10^{-6}$ | $1.12 \cdot 10^{-4}$ | $5.97 \cdot 10^{-4}$ | $3.30 \cdot 10^{-2}$ |
| Residual | $1.81 \cdot 10^{-3}$ | $1.50 \cdot 10^{-2}$ | $1.02 \cdot 10^{-1}$ | 1.86 |

Note: Trait heritabilities were estimated from the unnested models.

**Table A5:**
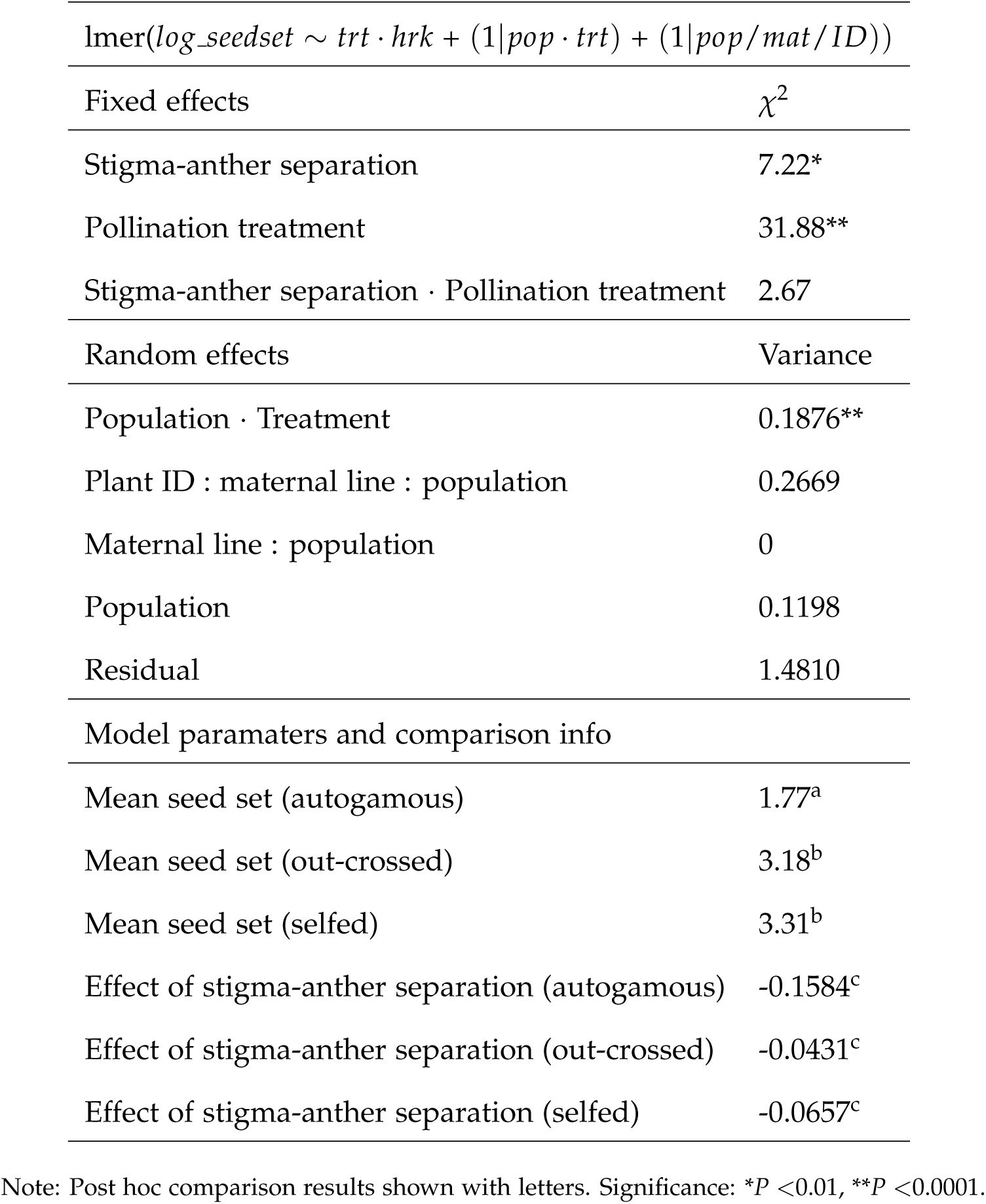
Model results of linear mixed-effects model comparing the effects of stigma-anther separation, pollination treatment and population on variation in seed set

**Table A6:** G matrix estimated using the subset of *Turnera ulmifolia* plants used in the pollination experiments

| | $Ela_m$ | EFN | FN | $S-A_d$ | Seed set |
| --- | --- | --- | --- | --- | --- |
| $Ela_m$ | 0.40 (0.22, 0.64) | | | | |
| EFN | 0.02 (−0.12, 0.16) | 0.25 (0.12, 0.44) |  |  |  |
| FN | −0.01 (−0.16, 0.15) | 0.06 (−0.08, 0.21) | 0.55 (0.35, 0.77) |  |  |
| $S-A_d$ | −0.07 (−0.20, 0.06) | 0.08 (−0.04, 0.21) | 0.09 (−0.04, 0.21) | 0.52 (0.38, 0.69) | |
| Seed set | **0.19 (0.06, 0.34)** | 0.03 (−0.09, 0.16) | 0.00 (−0.14, 0.16) | **−0.12 (−0.24, 0.00)** | 0.30 (0.15, 0.51) |

Note: We report the mean and 95 % HPD intervals of estimated genetic variances and co-variances. Values are bolded if significantly different from zero.

**Table A7:** Eigenanalysis of the G matrices fit for the five largest populations of *Turnera ulmifolia*

| (a) Brown's Town |  |  |  |  |
| --- | --- | --- | --- | --- |
| Eigenvalue | Elaiosome $m$ | EFN | FN | S-A separation |
| 0.8173 | -0.5196 | -0.1546 | 0.8121 | 0.2160 |
| 0.2244 | 0.7567 | 0.1855 | 0.3887 | 0.4918 |
| 0.1603 | 0.1709 | -0.9572 | -0.1254 | 0.1971 |
| 0.1351 | 0.3580 | -0.1595 | 0.4169 | -0.8201 |
| (b) Cave |  |  |  |  |
| Eigenvalue | Elaiosome $m$ | EFN | FN | S-A separation |
| 0.8435 | -0.2359 | -0.2870 | 0.2597 | -0.8914 |
| 0.3658 | 0.8803 | 0.3414 | 0.0850 | -0.3182 |
| 0.2014 | 0.4111 | -0.8946 | -0.0803 | 0.1559 |
| 0.1638 | 0.0203 | -0.0274 | 0.9586 | 0.2828 |
| (c) Mosquito Cove |  |  |  |  |
| Eigenvalue | Elaiosome $m$ | EFN | FN | S-A separation |
| 1.0008 | 0.2068 | -0.0980 | -0.5467 | -0.8055 |
| 0.3582 | 0.4739 | -0.6319 | -0.3967 | 0.4678 |
| 0.2546 | 0.6444 | 0.7235 | -0.1619 | 0.1873 |
| 0.2184 | 0.5634 | -0.2601 | 0.7194 | -0.3120 |
| (d) Murdock |  |  |  |  |
| Eigenvalue | Elaiosome $m$ | EFN | FN | S-A separation |
| 0.5400 | 0.8386 | 0.3975 | -0.0586 | 0.3678 |
| 0.2228 | 0.4054 | -0.8925 | -0.1975 | 0.0087 |
| 0.1610 | 0.0564 | -0.1837 | 0.9559 | 0.2223 |
| 0.1547 | -0.3594 | -0.1081 | -0.2095 | 0.9029 |
| (e) Saint Ann's Bay |  |  |  |  |
| Eigenvalue | Elaiosome $m$ | EFN | FN | S-A separation |
| 0.5968 | 0.8638 | 0.1701 | 0.0062 | -0.4741 |
| 0.1565 | -0.3806 | -0.3500 | -0.2385 | -0.8221 |
| 0.1075 | 0.3288 | -0.8770 | -0.2085 | 0.2816 |
| 0.0938 | 0.0291 | 0.2819 | -0.9485 | 0.1417 |

Note: The trait loadings for each measured phenotype are presented for each of four eigenvectors of each matrix.

**Table A8:** Eigendecomposition of the 4^th^ order genetic covariance tensor, Σ

| Eigenvalue of $\Sigma$ | Eigenvalue of E | Elaiosome $m$ | EFN | FN | S-A separation |
| --- | --- | --- | --- | --- | --- |
| 0.1507 | -0.9420 | 0.2267 | -0.0714 | -0.7568 | -0.6090 |
| 0.1507 | 0.2970 | 0.8818 | 0.4302 | 0.0835 | 0.1741 |
| 0.1507 | -0.1365 | -0.3460 | 0.8421 | 0.1324 | -0.3920 |
| 0.1507 | 0.0756 | -0.2265 | 0.3173 | -0.6347 | 0.6673 |
| 0.1130 | -0.9245 | 0.1171 | 0.2601 | -0.0745 | 0.9555 |
| 0.1130 | 0.3549 | -0.6199 | -0.2167 | 0.7294 | 0.1919 |
| 0.1130 | -0.1386 | -0.1388 | 0.9399 | 0.2196 | -0.2217 |
| 0.1130 | -0.0062 | 0.7634 | -0.0450 | 0.6436 | -0.0311 |

Note: Eigenvalues of $\Sigma$ describe axes of variation among G matrices and can be decomposed into eigentensors (E).
Each eigenvalue of E explains some amount of the variation captured by the eigenvalue of $\Sigma$ . We present the trait loadings of individual phenotypes on the eigentensors (E) of $\Sigma$ to quantify their contribution to differences among G.

**Table A9:** Phenotypic variance-covariance (P) matrices for the Jamaican metapopulation and five largest individual populations of *Turnera ulmifolia* we sampled

| (a) Jamaica |  |  |  |  |
| --- | --- | --- | --- | --- |
| | Elaiosome $m$ | EFN | FN | S-A separation |
| Elaiosome<sub>m</sub> | 1.00 |  |  |  |
| EFN | −0.07 (−0.12, −0.01) | 1.00 |  |  |
| FN | 0.04 (−0.01, 0.10) | **0.08 (0.02, 0.13)** | 1.00 |  |
| S-A sep. | −0.06 (−0.12, −0.01) | 0.05 (0.00, 0.11) | **0.13 (0.08, 0.18)** | 1.00 |
| (b) Brown's Town |  |  |  |  |
| | Elaiosome $m$ | EFN | FN | S-A separation |
| Elaiosome<sub>m</sub> | 1.00 |  |  |  |
| EFN | 0.11 (−0.10, 0.31) | 1.00 |  |  |
| FN | −0.07 (−0.26, 0.13) | 0.11 (−0.10, 0.32) | 1.00 |  |
| S-A sep. | 0.19 (−0.02, 0.39) | 0.07 (−0.12, 0.25) | −0.07 (−0.22, 0.09) | 1.00 |
| (c) Cave |  |  |  |  |
| | Elaiosome $m$ | EFN | FN | S-A separation |
| Elaiosome<sub>m</sub> | 1.00 |  |  |  |
| EFN | 0.02 (−0.15, 0.20) | 1.00 |  |  |
| FN | −0.07 (−0.29, 0.15) | 0.13 (−0.11, 0.35) | 1.00 |  |
| S-A sep. | 0.08 (−0.15, 0.31) | −0.07 (−0.27, 0.13) | 0.07 (−0.15, 0.28) | 1.00 |
| (d) Mosquito Cove |  |  |  |  |
| | Elaiosome $m$ | EFN | FN | S-A separation |
| Elaiosome<sub>m</sub> | 1.00 |  |  |  |
| EFN | 0.01 (−0.18, 0.20) | 1.00 |  |  |
| FN | −0.04 (−0.25, 0.18) | −0.11 (−0.32, 0.12) | 1.00 |  |
| S-A sep. | 0.02 (−0.19, 0.22) | −0.01 (−0.18, 0.16) | 0.22 (−0.01, 0.42) | 1.00 |
| (e) Murdock |  |  |  |  |
| | Elaiosome $m$ | EFN | FN | S-A separation |
| Elaiosome<sub>m</sub> | 1.00 |  |  |  |
| EFN | 0.14 (−0.05, 0.33) | 1.00 |  |  |
| FN | 0.15 (−0.05, 0.34) | **0.21 (0.00, 0.40)** | 1.00 |  |
| S-A sep. | **0.27 (0.06, 0.46)** | 0.01 (−0.17, 0.19) | 0.01 (−0.15, 0.17) | 1.00 |
| (f) Saint Ann's Bay |  |  |  |  |
| | Elaiosome $m$ | EFN | FN | S-A separation |
| Elaiosome<sub>m</sub> | 1.00 |  |  |  |
| EFN | 0.20 (−0.01, 0.40) | 1.00 |  |  |
| FN | 0.11 (−0.10, 0.31) | −0.07 (−0.26, 0.13) | 1.00 |  |
| S-A sep. | −0.05 (−0.23, 0.13) | −0.10 (−0.29, 0.11) | −0.01 (−0.17, 0.15) | 1.00 |

Note: We estimated mean and 95% confidence intervals for phenotypic correlations. Estimates significantly different than zero are bolded.

**Table A10:** Phenotypic divergence variance-covariance matrix of the population means for mutualistic traits (D) in *Turnera umifolia*

| | $Ela_m$ | EFN | FN | S-$A_d$ |
| --- | --- | --- | --- | --- |
| $Ela_m$ | **0.54 (0.26, 1.08)** | | | |
| EFN | 0.01 (−0.17, 0.19) | 0.20 (0.09, 0.39) |  |  |
| FN | 0.03 (−0.26, 0.34) | 0.10 (−0.05, 0.32) | **0.51 (0.25, 1.04)** |  |
| S-$A_d$ | −0.11 (−0.50, 0.21) | 0.05 (−0.14, 0.28) | 0.13 (−0.16, 0.52) | **0.66 (0.32, 1.35)** |

Note: We report the mean and 95 % HPD intervals of estimated variances and co-variances among population means. Estimates significantly different than zero are bolded.

**Table A11:** Genetic variance-covariance matrix of the island-wide metapopulation of *Turnera umifolia*. Phenotypic trait values were standardized by mean but not to unit variance

| | $Ela_m$ | EFN | FN | S-$A_d$ |
| --- | --- | --- | --- | --- |
| $Ela_m$ | 0.035 (0.029, 0.042) | | | |
| EFN | −0.0066 (−0.0019, 0.0016) | 0.081 (0.060, 0.11) |  |  |
| FN | 0.0058 (−0.0019, 0.0016) | **0.052 (0.021, 0.085)** | 0.34 (0.27, 0.42) |  |
| S-$A_d$ | −0.0055 (−0.0022, 0.011) | **0.059 (0.023, 0.096)** | **0.10 (0.041, 0.16)** | 0.51 (0.42, 0.60) |

Note: We report the mean and 95 % HPD intervals of estimated variances and co-variances among population means. Estimates of genetic co-variances significantly different from zero are bolded. We did not test the statistical significance of genetic variances (i.e., diagonal elements) here, but see Table A3

## Online figure legends

**Figure A1:**
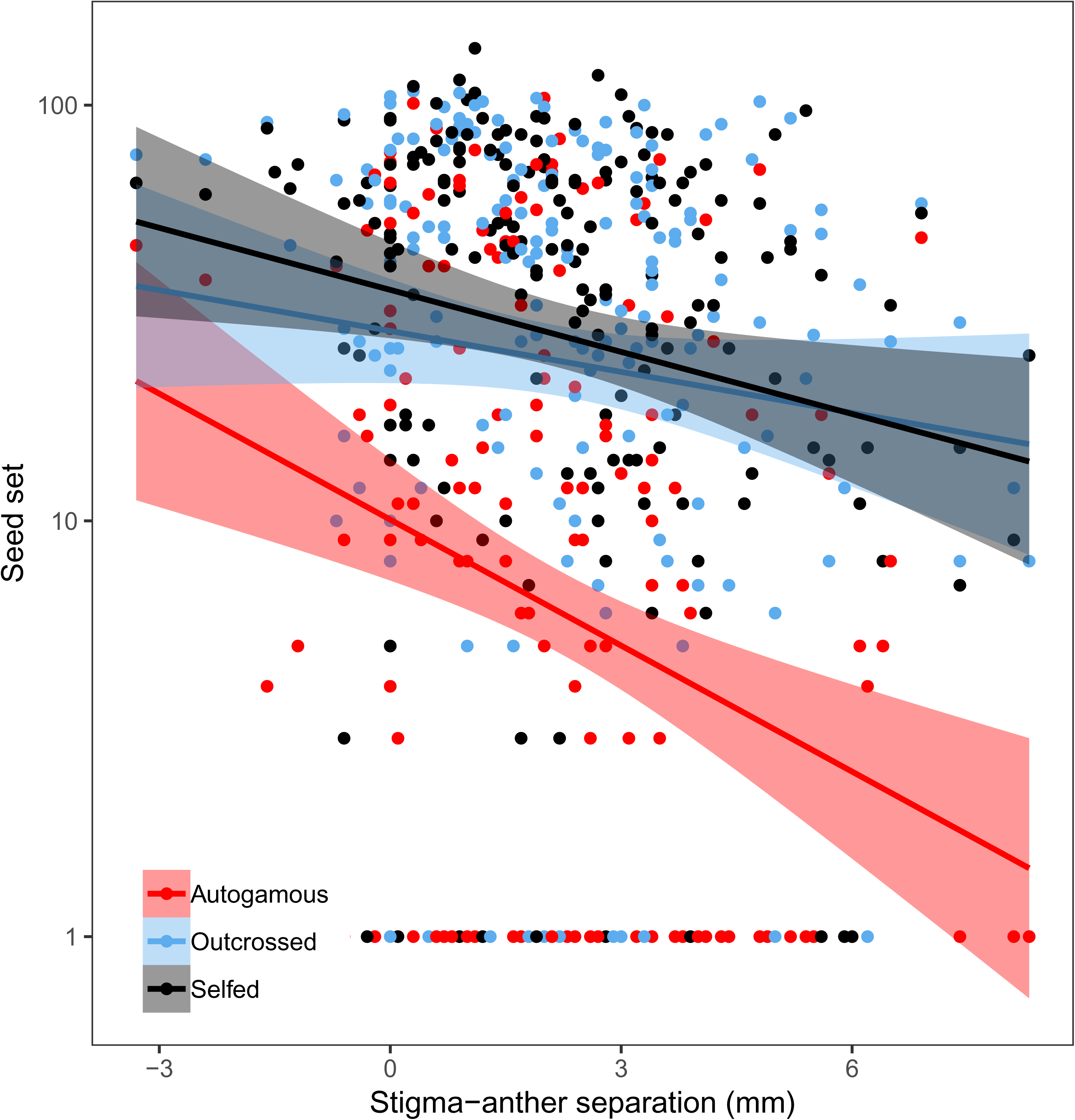
Seed set of pollinated *Turnera ulmifolia* flowers regressed against stigma-anther separation. We pollinated stigmas with either pollen from the same flower (selfed), from a different plant of the same population (out-crossed), or allowed pollination to occur upon withering of the perianth (autogamous).

**Figure A2:**
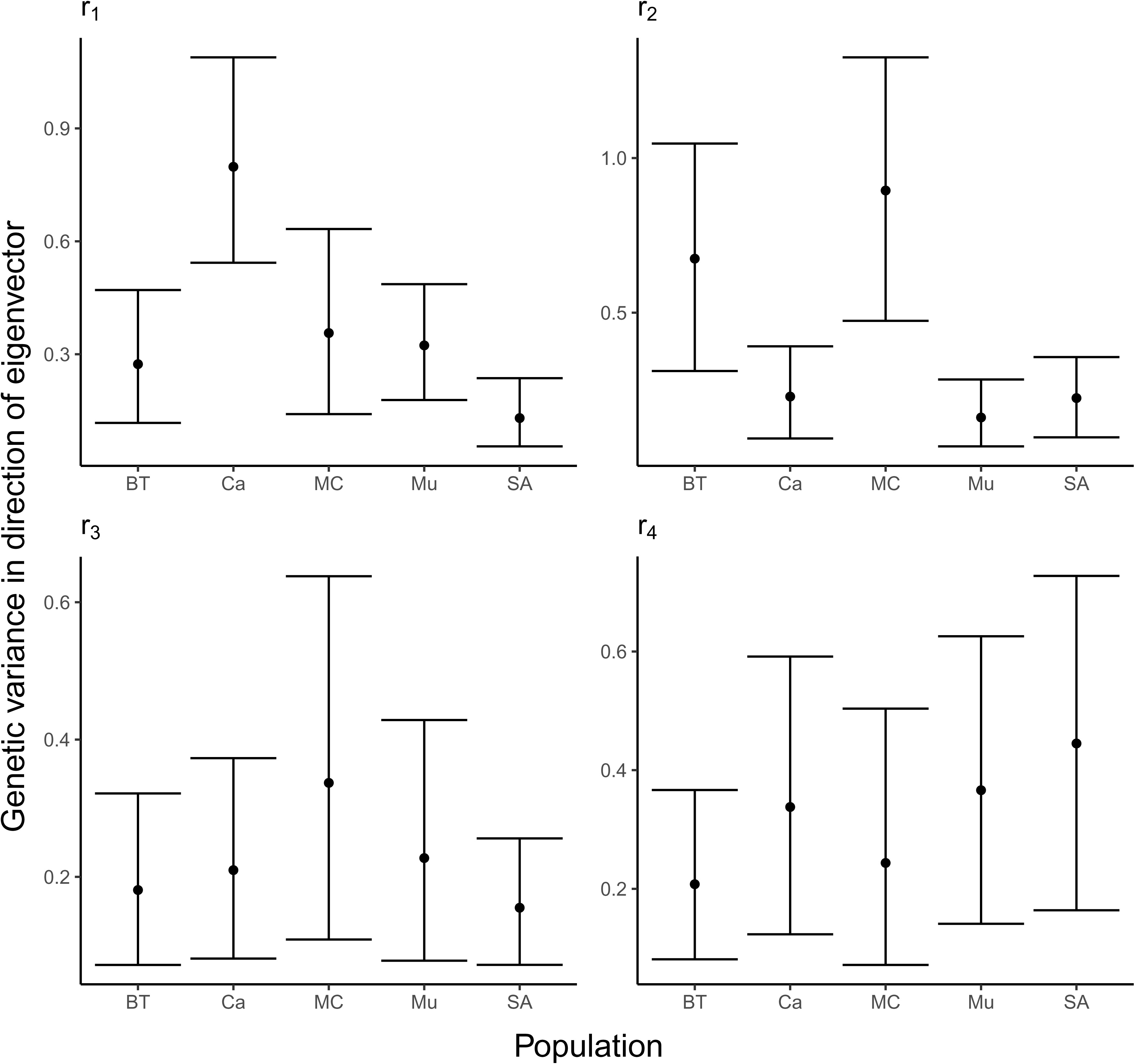
Results from the implementation of modified random skewers. Genetic variance in the direction of the four eigenvectors (r_1_ – r_4_) of R for the 5 largest Jamaican populations of *Turnera ulmifolia* we sampled (Brown’s Town (BT), Cave (Ca), Mosquito Cove (MC), Murdock (Mu), and Saint Ann’s Bay (SA)). All error bars are 95% HPD intervals.

**Figure A3:**
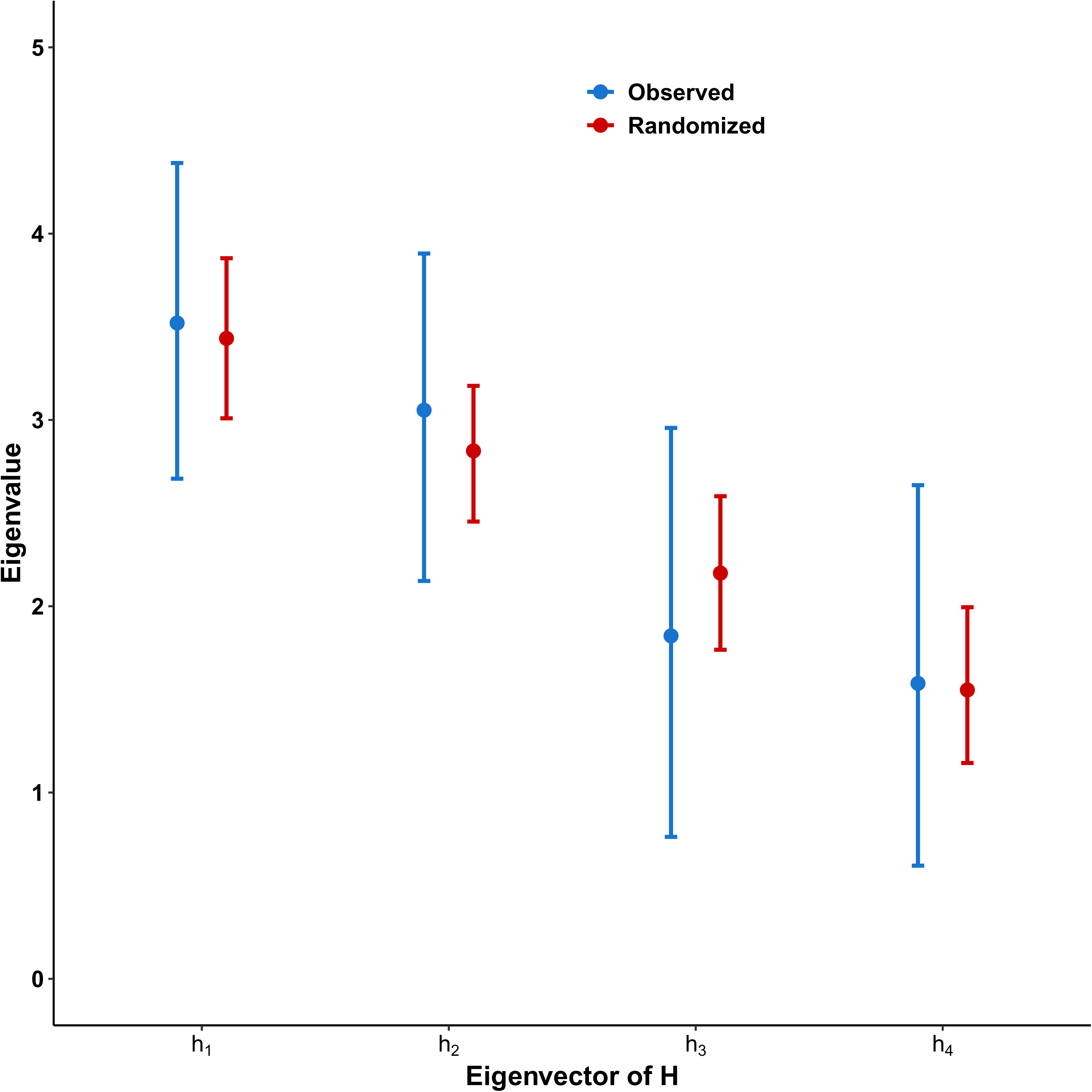
Results from the Krzanowski common subspace analysis, showing the eigenvalues of H for observed (blue) and randomized (red) G matrices. The analysis was performed with the number of eigenvectors of the summary matrix H set to 2 (half of the total traits). All error bars are 95% HPD intervals.

**Figure A4:**
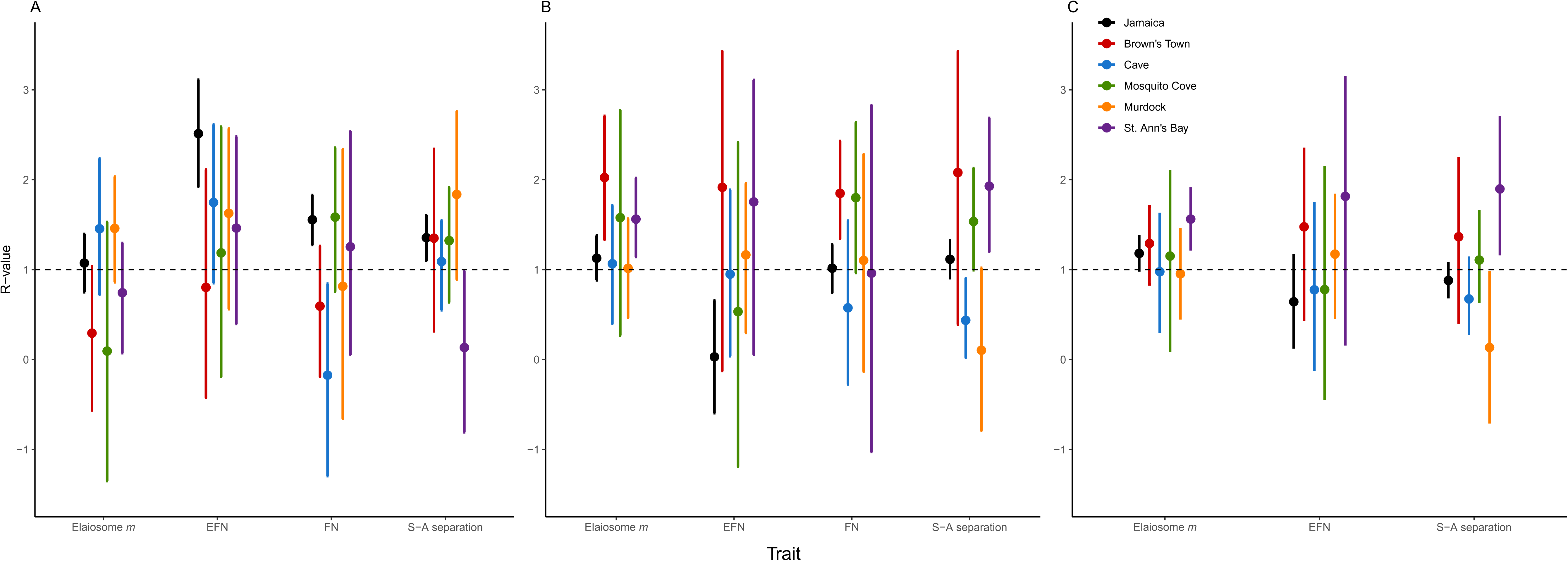
R values with 95% HPD intervals for the Jamaican metapopulation and our five largest *Turnera ulmifolia* populations. Values greater than 1 (dashed line) indicate that genetic correlations among traits facilitate responses to selection, while values less than 1 indicate that genetic correlations constrain their responses. R values were calculated under selection scenarios where (a) all traits experienced positive directional selection (*β* = 0.1), (b) where elaiosome mass and EFN are under positive directional selection while FN and stigma-anther separation are under negative directional selection (*β* = -0.1), and (c) where elaiosome mass and EFN are under positive selection, stigma-anther separation is selected against, and FN is selectively neutral (*β* = 0).

